# Activation of Notch1 drives the development of radiation-induced thymic lymphoma in p53 wild-type mice

**DOI:** 10.1101/2020.08.21.261487

**Authors:** Chang-Lung Lee, Kennedy Davis Brock, Stephanie Hasapis, Dadong Zhang, Alexander B. Sibley, Xiaodi Qin, Jeremy Gresham, Isibel Caraballo, Lixia Luo, Andrea R. Daniel, Matthew J. Hilton, Kouros Owzar, David G. Kirsch

## Abstract

Mouse models of radiation-induced thymic lymphoma are widely used to study the development of radiation-induced blood cancers and to gain insights into the biology of human T-lymphoblastic leukemia/lymphoma. Here, we aimed to determine key oncogenic drivers for the development of radiation-induced thymic lymphoma by performing whole-exome sequencing using tumors and paired normal tissues from mice with and without irradiation. Thymic lymphomas from irradiated wild-type (WT), p53^+/-^ and Kras^LA1^ mice were not observed to harbor significantly higher numbers of non-synonymous somatic mutations compared to thymic lymphomas from unirradiated p53^-/-^ mice. However, we observed distinct patterns of recurrent mutations in genes that control the Notch1 signaling pathway based on the mutational status of p53. Preferential activation of Notch1 signaling in p53 WT lymphomas was also observed at the RNA and protein level. Using reporter mice for activation of the Notch1 signaling we observed that TBI enriched Notch1^hi^ CD44^+^ thymocytes, which are capable of self-renewal *in vivo*. Mechanistically, genetic inhibition of Notch1 signaling in thymocytes prevented the formation of radiation-induced thymic lymphoma in p53 WT mice. Taken together, our results demonstrate a critical role of activated Notch1 signaling in driving multi-step carcinogenesis of thymic lymphoma following total-body irradiation in p53 WT mice.

## INTRODUCTION

Exposure of a significant part of the bone marrow to ionizing radiation is associated with an excess risk of developing blood cancers, including acute myeloid leukemia and acute lymphoblastic leukemia^1–4^. Experimentally, one of the widely used mouse models to study the carcinogenic effect of total-body irradiation (TBI) on hematopoietic cells is radiation-induced thymic lymphoma^5^. Dr. Henry Kaplan and colleagues first reported that TBI of C57 black mice leads to thymic lymphomas arising from T-cell precursors in the thymus^6,7^. Lymphoma cells that develop in the thymus can become leukemic and disseminate to other organs including the spleen, liver, kidney and bone marrow^8^. Intriguingly, the incidence of radiation-induced thymic lymphoma after TBI depends on the daily radiation dose, number of fractions, time interval between each fraction, and the accumulated total radiation dose^7^. For example, exposure of C57BL/6 mice to 1.6 to 1.8 Gy TBI once per week for 4 consecutive weeks causes more than 90% of mice to develop thymic lymphoma within 8 months after irradiation. Even though this classic model has been utilized by many laboratories over the past 70 years, mechanisms by which TBI induces thymic lymphoma in wild-type mice are incompletely understood.

The application of next-generation sequencing provides an opportunity to gain novel insights into mechanisms of radiation-induced carcinogenesis. For example, whole-exome sequencing (WES) studies of mouse solid tumors uncovered mutational signatures enriched in murine radiation-induced solid tumors compared to radiation naïve tumors^9,10^. In addition, genomic analyses of human cancers reveal specific patterns of single-nucleotide substitutions and DNA deletions among radiation-associated solid cancers^11,12^. Moreover, results from WES and whole-genome sequencing show distinct mutational patterns in radiation-induced cancers that develop in wild-type mice versus mice with pre-existing germline mutations. For example, using a mouse model of B-cell lymphoma, Patel et al. found that gamma-rays and heavy ion (^56^Fe) radiation markedly increased the number of insertions and deletions (indels) per tumor only in mice lacking one allele of the mismatch repair gene *Mlh1* (Mlh1^+/-^)^13^. Also, using mice carrying different loss of function alleles of the tumor suppressor p53, Li et al. observed that the genomic landscape of radiation-induced cancers is influenced by radiation quality, germline *p53* deficiency and tissue/cell of origin^14^. Collectively, findings from these genomic studies advance our understanding of mutational processes that underlie the development of radiation-induced cancers. However, although these next-generation sequencing studies have identified mutations in many putative driver genes, critical oncogenic pathways that are necessary to drive the development of radiation-induced cancers remain undefined.

Here, we performed WES to define somatic mutations and copy number alterations of coding genes in thymic lymphoma that develop in mice following 4 weekly fractions of 1.8 Gy TBI. We conducted our study not only in wild-type (C57BL/6J) mice, but also in genetically engineered mice predisposed to thymic lymphomas including Kras^LA1^ mice^15^ and p53^+/-^ mice^16,17^. As a control, we also analyzed thymic lymphomas in unirradiated mice that developed as a result of germline deletion of both *p53* alleles in hematopoietic cells in order to compare the mutational signatures between thymic lymphomas that develop in irradiated mice versus radiation naïve mice. Importantly, utilizing the discoveries from whole-exome sequencing, we performed mechanistic studies to identify Notch1 signaling as a critical oncogenic pathway that drives the formation of radiation-induced thymic lymphoma in wild-type mice.

## MATERIALS AND METHODS

### Mouse strains and induction of thymic lymphomas

All animal procedures for this study were approved by the Institutional Animal Care and Use Committee (IACUC) at Duke University. Mouse strains used to generate thymic lymphomas with and without irradiation were described previously including wild-type C57BL/6J mice (the Jackson Laboratory #000664), B6.SJL mice (the Jackson Laboratory #002014), Kras^LA1^ mice on a B6 background^15,18^, p53^+/-^ mice^8,16^, Tie2Cre; p53^FL/-^ mice^19^, LckCre mice^20^ (the Jackson Laboratory #003802), Rbpj floxed mice^21^ and Rbpj-Venus reporter mice^22^ (the Jackson Laboratory #020942). To study radiation-induced thymic lymphomas, mice were exposed to 1.8 Gy total-body irradiation (TBI) every week for 4 consecutive weeks (1.8 Gy x 4)^8^. The irradiation experiments were performed using the X-RAD 320 Biological Irradiator (Precision X-ray). Mice were irradiated at 50 cm from the radiation source with a dose rate of 3.18 Gy/min with 320 kVp X-rays, using 12.5 mA and a filter consisting of 2.5 mm Al and 0.1 mm Cu. The dose rate was measured with an ion chamber by members of the Radiation Safety Division at Duke University.

### Whole exome sequencing (WES)

To prepare samples for WES, thymic lymphoma specimens stored in RNAlater (Invitrogen) and matched tail that were snap frozen were used for DNA extraction. DNA extraction was performed using the DNeasy Blood and Tissue Kit or the AllPrep DNA/RNA Mini Kit (Qiagen) per the manufacturer’s guidelines. WES was performed using previously described methods^9^. Sequencing libraries were prepared using the Agilent SureSelect^*XT*^ Mouse All Exon Kit (S0276129). The kit has a target size of 49.6 megabases. Methods for data analyses including somatic mutation calling, generation of somatic mutation plots, generation of mutational signatures, and assessment of copy number variations (CNVs) were described previously^9^, with two exceptions. First, in this study, genes with absolute value of estimated CNVs greater than 0.3 were considered CNVs. Second, for the purposes of inclusion in the heat maps, mean CNV was calculated across all samples, and oncogenes with mean CNV greater than zero and tumor suppressor genes with mean CNV less than zero were included. The sequencing data along with the called mutations in vcf format have been deposited into the National Center for Biotechnology Information Sequence Read Archive under project ID PRJNA627412.

### Notch1 Exon 1 Deletion PCR

DNA was extracted from thymic tissues using the *Quick-DNA* Miniprep Plus Kit (Zymo Research) according to the manufacturer’s protocols. PCR was performed using Phusion polymerase (New England Biolabs) using a 90 second elongation for 35 cycles. Primers used were: Notch1 1Δ F 5’-ATGGTGGAATGCCTACTTTGTA-3’, Notch1 1Δ R 5’-CGTTTGGGTAGAAGAGATGCTTTAC-3’ (Annealing Temp = 62.8°C, product size ~500 bp), Notch1 E30E31 F 5’-CACATGCACACATTCCCTAGC-3’, Notch1 E30E31 R 5’-GCTTTGCAGCATCTGAACGA-3’ (Annealing Temp = 64.4°C, Product size ~1000 bp).

### qRT-PCR to detect mRNA expression

RNA was isolated from thymic tissues using Direct-zol RNA Miniprep Kit (Zymo Research) according to the manufacturer’s protocols. cDNA was synthesized using iScript cDNA Synthesis Kit (Bio-Rad). Gene expression for Hes1(Mm01342805_m1), Ikzf1(Mm01187878_m1), Notch1 5’(Mm00627192_m1), and Notch1 3’(Mm01326754_g1) were analyzed using Taqman probes and TaqMan™ Universal PCR Master Mix (ThermoFischer) with Gapdh(Mm99999915_g1) as an internal control. Gene expression for Dtx1 (qMmuCID0027483) and Myc (qMmuCID0006528) were analyzed using PrimePCR SYBR Green Assays and SsoAdvanced Universal SYBR Green Supermix (BioRad) and with TBP (Fwd: 5’-TGCACAGGAGCCAAGAGTGAA-3’, Rev: 5’-CACATCACAGCTCCCCACCA-3’) as an internal control. Samples were run in triplicate on the QuantStudio 6 Flex Real-Time PCR System.

### qRT-PCR to detect DNA copy numbers

Copy number variants determined by WES for NF2 and Ikzf1 were tested using the same genomic DNA. Gene expression was measured using PowerUp SYBR Green Master Mix (Thermo Fisher Scientific) in the QuantStudio 6 Flex Real-Time PCR System in triplicate with 10 ng of gDNA per well. We utilized the following primer sequences for the target genes: NF2E12 (Fwd: 5’-GAGGAGGAGGCCAAGCTTTT-3’, Rev: 5’-TCCTCTCTGACTCCTCAGCC-3’); Ikzf1E5 (Fwd: 5’-AATGAATGGCTCCCACAGGG-3’, Rev: 5’-GAACCATGAGCACATTGGGC-3’). qRT-PCR expression results from lymphoma samples were normalized using ΔΔC_t_ analysis, with the TFRC gene and paired tail samples as internal controls. The TFRC primer sequence (Fwd: 5’–TGTAATTGGGTATAGGCTTGCA-3’, Rev: 5’–ATCCTGTACTCAAGAATTGGCT-3’) has been previously described^24^.

### Immunoblotting

Proteins were extracted from cells or tissues using PARIS kit (Invitrogen). The protein concentration was determined using Pierce Rapid Gold BCA Protein Assay (Thermo Scientific) according to the manufacturer’s protocol. Approximately 20 mg of total protein were loaded for electrophoresis into 4–20% sodium dodecyl sulfate polyacrylamide gels (Bio-Rad). Separated proteins were transferred to a 0.45 μm nitrocellulose membrane (Bio-Rad). Membranes were blocked with Odyssey Blocking Buffer (Li-Cor) diluted 1:1 with tris buffered saline (TBS). Protein levels were detected using antibodies against Intracellular Domain of Notch1 (ICN) (Val1744 Cell Signaling Technology #4147, D3B8, 1:500 dilution) and β-actin (Millipore-Sigma, A5441, clone AC-15, 1:5,000 dilution) followed by secondary fluorescently-conjugated IRDye 800CW and IRDye 680RD antibodies (Li-Cor, 1:15,000 dilution). Bands were visualized at 700 and 800nm using the Odyssey CLX imaging system (Li-Cor).

### Validation of SNVs using Sanger sequencing

Lymphoma single nucleotide variants (SNVs) of Notch1, Ikzf1, Pten, Akt1, and p53 were determined by WES were validated by Sanger sequencing. The sequences sent for Sanger sequencing were amplified by PCR using AccuPrime Taq DNA Polymerase (Thermo Fisher Scientific) and performed on the same genomic DNA sent for WES. Primer sequences are described in **Table S1**. The PCR Protocol for Notch1 exon 34 primers 3 through 5 and Ikzf1 exon 5 was: denaturing stage of 94°C for 2 minutes; 40 cycles of 94°C for 30 seconds, annealing temperature for 30 seconds, and 68°C for 60 seconds; followed by 4°C forever. Notch1 exon 34 primers 1 and 2 differed only in extending the elongation step of 68°C, for 90 seconds instead of 60 seconds. The PCR protocols for Notch1 exons 27 and 26, Ikzf1 exons 6, 8, and 9, TRP53 exon 7, Pten exons 7 and 8, and Akt1 exon 4 utilized the same protocol except for the elongation step, which was 45 seconds. Sequencing and mutation detection were performed by Eton Bioscience, Inc.

### Flow cytometry

Total thymocytes were isolated from the thymus in HSC buffer (HBSS with Ca2^+^ and Mg2^+^, 5% fetal bovine serum, 2 mM EDTA). Red blood cells (RBC) were lysed using ACK lysing buffer (Lonza). 1 x 10^6^ live thymocytes were blocked with a rat anti-mouse CD16/32 antibody (BD Pharmingen) and stained with a combination of the antibodies including PE-Cy5 conjugated anti-mouse CD4 (clone: GK1.5, eBioscience), PE conjugated anti-mouse CD8 (clone: 53-6.7, eBioscience), APC conjugated anti-mouse CD44 (clone: IM7, eBioscience), BV421 conjugated anti-mouse CD25 antibodies (clone: PC61, BioLegend). FITC conjugated anti-mouse CD45.2 (clone: 104, eBioscience), and APC-eFluor780 conjugated anti-mouse CD45.1 (clone: A20, e eBioscience). All antibodies were diluted 1:400. Data were collected from at least 100,000 single cells by FACSCanto (BD Pharmingen) and analyzed by FlowJo (Tree Star, Inc) without knowledge of the genotype or treatment by a single observer (C-LL).

### Thymocyte transplantation

Thymocyte transplantation was performed according to a method described previously^23^ with slight modifications. Individual recipient mice were exposed to 5 Gy TBI and then received 2 x 10^7^ thymocytes from donor mice through intravenous injection approximately 6 hours after irradiation. Twenty-one days after thymocyte transplantation, thymocytes were harvested from recipient mice for subsequent analysis using flow cytometry.

### Statistics

Statistical results presented here were post-hoc analyses. The p-values reported were two-sided and were not adjusted for multiple testing. Pairwise comparisons of quantitative phenotypes were based on the Mann-Whitney U test. Comparisons of three or more groups used the Kruskal-Wallis test. Comparisons of proportions were based on the chi-squared test. Time-to-event analyses were based on the Cox proportional hazards model and reported p-values are from the score test. Kendall’s W was used to test concordance of estimated CNV by qPCR and WES. Inferential analyses were conducted using the R statistical environment^24^ and extension packages from the Comprehensive R Archive Network (https://cran.r-project.org/) and the Bioconductor project^25^. Box-and-whisker plots presented in the figures were constructed as previously described^9^. For all experiments, data are presented as mean ± SE and each data point represents one mouse or tumor sample.

The knitr package^26^ was used to generate dynamic reports and conduct the analyses in accordance with the principles of reproducible analysis. Duke’s GitLab was used for source code management (https://gitlab.oit.duke.edu). The code for replicating the statistical analyses presented here has been made available through a public source code repository (https://gitlab.oit.duke.edu/dcibioinformatics/pubs/kirsch-lee_lymphoma).

## RESULTS

### Somatic mutation analysis of thymic lymphomas from irradiated and unirradiated mice

Thymic lymphomas were generated in irradiated and unirradiated mice on four distinct genotypes (**Figure S1**). Radiation-induced thymic lymphomas were generated in wild-type (WT) (C57BL/6J) mice, Kras^LA1^ mice^15^ or p53^+/-^ mice^16,17^ that were exposed to 1.8 Gy total-body irradiation (TBI) per week for 4 consecutive weeks^7,8^. Unirradiated Tie2Cre; p53^FL/-^ mice (Non-IR p53^-/-^) in which both alleles of the p53 gene are deleted in endothelial cells and hematopoietic cells^19^ were used to generate thymic lymphomas in the absence of irradiation. The mean time for the formation of radiation-induced thymic lymphoma in WT mice was approximately 150 days after irradiation. The latency of thymic lymphomas in Kras^LA1^ mice and p53^+/-^ mice was reduced to approximately 100 days post-irradiation because these genetically engineered mice are predisposed to develop thymic lymphoma^15,17^. The latency of radiation-induced thymic lymphomas is comparable to time of spontaneous thymic lymphoma development in Non-IR p53^-/-^ mice (**Figure S1**).

Thymic lymphomas and paired normal tissues were subjected to WES to identify somatic mutations that occurred specifically in tumors (**Figure 1**). This analysis showed that while the number of total somatic mutations per tumor differed between thymic lymphomas induced by TBI alone, TBI plus genetic predisposition (Kras^LA1^ and p53^+/-^), and the loss of both alleles of p53 in the absence of TBI (Kruskal-Wallace p-value= 0.0221), the number of nonsynonymous somatic mutations was not significantly different among different lymphoma groups (Kruskal-Wallace p-value=0.0677, **Figure 1, a and b**). Notably, we observed that two radiation-induced lymphomas, tumor numbers 5015 (WT) and 5020 (Kras^LA1^), exhibited substantially higher numbers of somatic mutations compared to other lymphomas (**Figure 1a**). Examination of the proportion of indels and SNVs did not show statistically significant difference among different lymphoma groups (Kruskal-Wallace p-value=0.9099, **Figure 1, c to e and Figure S2**). In addition, unsupervised hierarchical clustering of thymic lymphomas based on data from SNVs did not differentiate radiation-induced thymic lymphomas from spontaneous tumors that arose in Non-IR p53^-/-^ mice (**Figure 1f**). However, this analysis revealed a distinct cluster comprised tumor numbers 5015 and 5020, two radiation-induced thymic lymphomas with the highest number of somatic mutations (**Figure 1a**).

**Figure 1.**
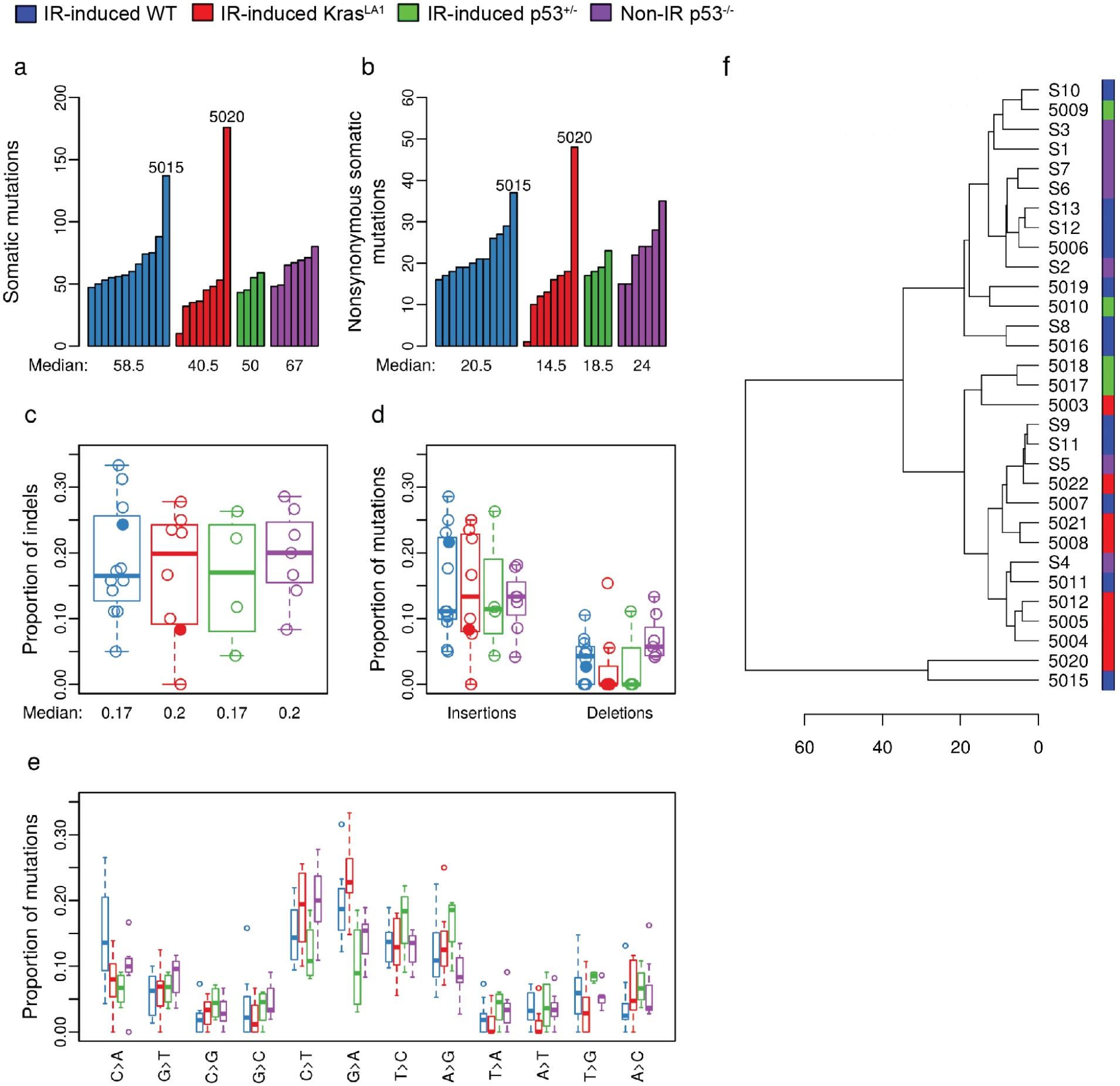
Somatic mutations in murine thymic lymphomas. Thymic lymphomas were harvested from the following cohorts of mice: IR-induced WT (blue), IR-induced Kras^LA1^ (red), IR-induced p53^+/-^ (green) and Non-IR p53^-/-^ (purple). **a,** The number of total somatic mutations per tumor. **b,** The number of somatic nonsynonymous mutations per tumor. **c,** The proportion of insertion-deletions (indels) within nonsynonymous mutations. **d,** The proportions of insertions and deletions within nonsynonymous mutations. **e,** The proportions of different single nucleotide variants. **f,** Unsupervised hierarchical clustering of thymic lymphomas based on data from single nucleotide variants. All panels illustrate the data for n=31 tumors. Data from lymphomas that had substantially higher number of somatic mutations (5015 and 5020) were labeled or denoted by closed circles.

To generate mutational signatures common to each lymphoma genotype, we performed a signature analysis as described by Alexandrov, et al.^27^ and compared mutational signatures of mouse lymphomas with the Catalogue Of Somatic Mutations In Cancer (COSMIC) mutational signatures of human cancers (https://cancer.sanger.ac.uk/cosmic/signatures) (**Figure S3**). Among radiation-induced thymic lymphomas, all three genotypes exhibited cosine-similarity with three groups of COSMIC mutational signatures of human cancers (**Figure S3, a to c**): defective DNA mismatch repair (Signatures 6, 15 and 26), spontaneous deamination of 5-methylcytosine (Signature 1) and a signature with unknown etiology that has been found in all cancer types (Signature 5). Remarkably, the same groups of signatures were also found in Non-IR p53^-/-^ thymic lymphomas (**Figure S3d**). Together, as shown in the analysis of somatic mutations, our results did not reveal unique mutational processes underlying the development of thymic lymphomas induced by TBI alone, TBI plus genetic predisposition (Kras^LA1^ and p53^+/-^) or the loss of functional p53.

### Copy number variations analysis of murine thymic lymphomas

WES data was evaluated using CODEX2 for CNVs^28^. Somatic deletions and amplifications were detected across all 19 mouse chromosomes in the thymic lymphomas (**Figure 2a**). Among the four lymphoma cohorts, IR-induced Kras^LA1^ and IR-induced WT lymphomas had a generally low number of genes affected by CNVs. Interestingly, tumor numbers 5015 and 5020, which exhibited a higher number of somatic mutations compared to other lymphomas, also had the highest numbers of CNVs compared to other tumors within the same cohort (**Figure 2, b to d**). In contrast, the number of genes with CNVs was significantly higher in IR-induced p53^+/-^ lymphomas compared to IR-induced Kras^LA1^ and IR-induced WT lymphomas (pairwise Mann-Whitney U p-values 0.0040 and 0.0011, respectively, **Figure 2b**). While lymphomas with the greatest number of genes affected by CNVs were initiated in p53^+/-^ mice by TBI with loss of p53 function, the number of genes with CNVs was also significantly higher in Non-IR p53^-/-^ lymphomas compared to IR-induced Kras^LA1^ and IR-induced WT lymphomas (pairwise Mann-Whitney U p-values 0.0140 and 0.0283, respectively, **Figure 2b**). In sum, our results suggest that in mouse models of thymic lymphomas, TBI cooperates with loss of p53 to increase the number of CNVs.

**Figure 2.**
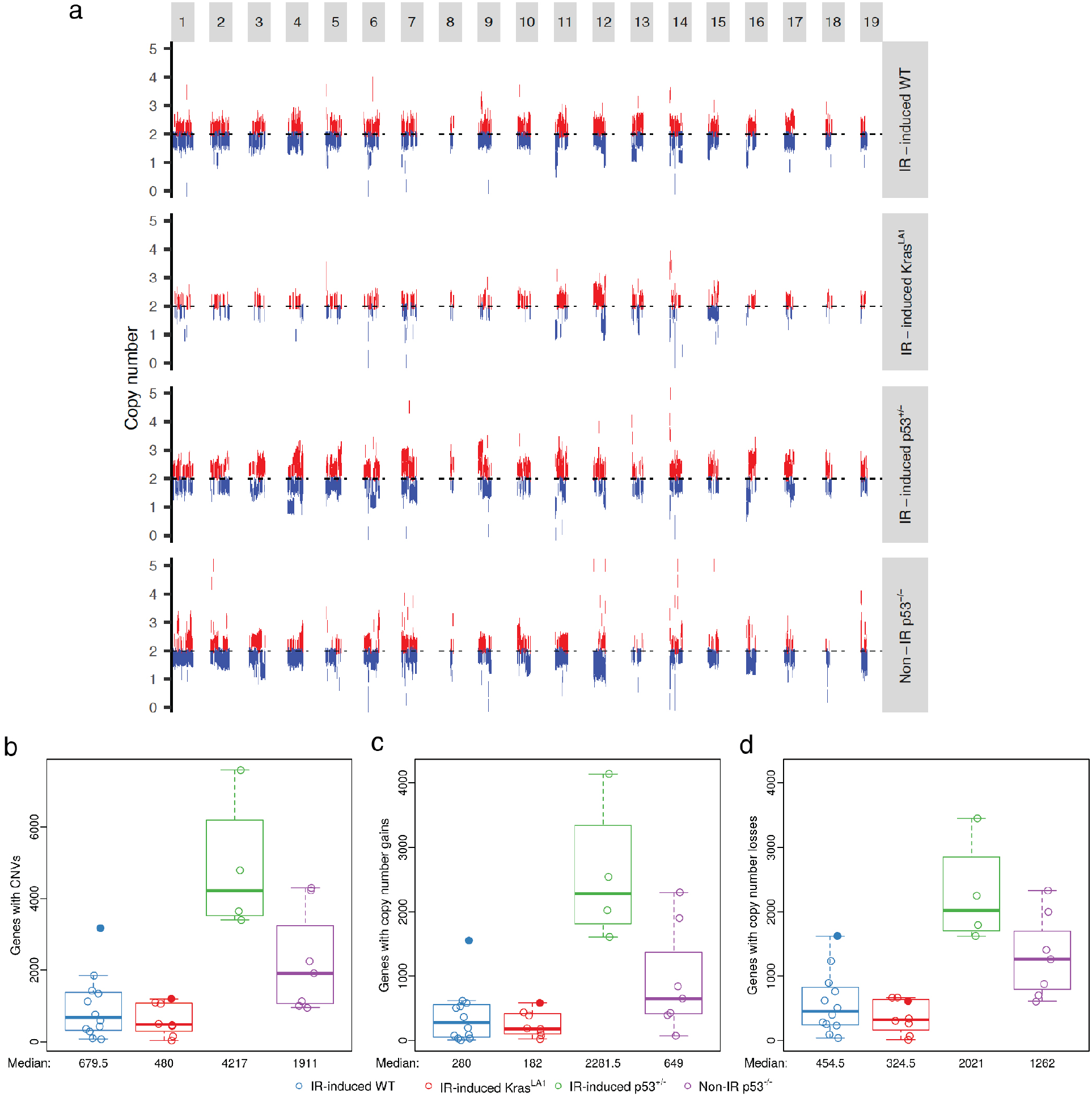
Somatic copy number variations (CNVs) in thymic lymphomas. **a,** Schematic of CNVs across the 19 mouse chromosomes. Results represent pooled data from lymphomas of the same cohort. DNA deletions and amplifications are labeled by blue and red, respectively. **b,** The number of genes affected by CNVs. **c,** The number of genes with copy number gains. **d,** The number of genes with copy number losses. Panels illustrate the data for n=31 tumors. Data from lymphomas that had substantially higher mutational loads (5015 and 5020) were denoted with closed circles.

### Identification of putative genetic drivers underlying T-lymphomagenesis

To investigate genetic alterations that potentially drive oncogenesis of thymic lymphomas, we identified genes from the COSMIC database that were impacted by somatic mutations and CNVs. In IR-induced WT lymphomas, nonsynonymous mutations were found in 1 to 5 COSMIC genes (median = 3) per tumor (**Figure 3a**). Even fewer nonsynonymous mutations in COSMIC genes were found in thymic lymphomas from mice that harbored a germline tumor-initiating mutation including irradiated Kras^LA1^ mice, irradiated p53^+/-^ mice, as well as radiation naïve p53^-/-^ mice (**Figure 3a**). The number of COSMIC genes altered by CNVs in radiation-induced lymphomas from either WT mice or Kras^LA1^ mice was significantly lower compared to radiation-induced lymphomas in p53^+/-^ mice (Kruskal-Wallace p-value = 0.0010, **Figure 3b)**. The tumors from unirradiated p53^-/-^ mice exhibited an intermediate level of CNVs in COSMIC genes (**Figure 3b**). Of note, tumor numbers 5015 and 5020 also had the highest number of COSMIC genes affected by CNVs compared to other tumors within the same cohort (**Figure 3b**).

**Figure 3.**
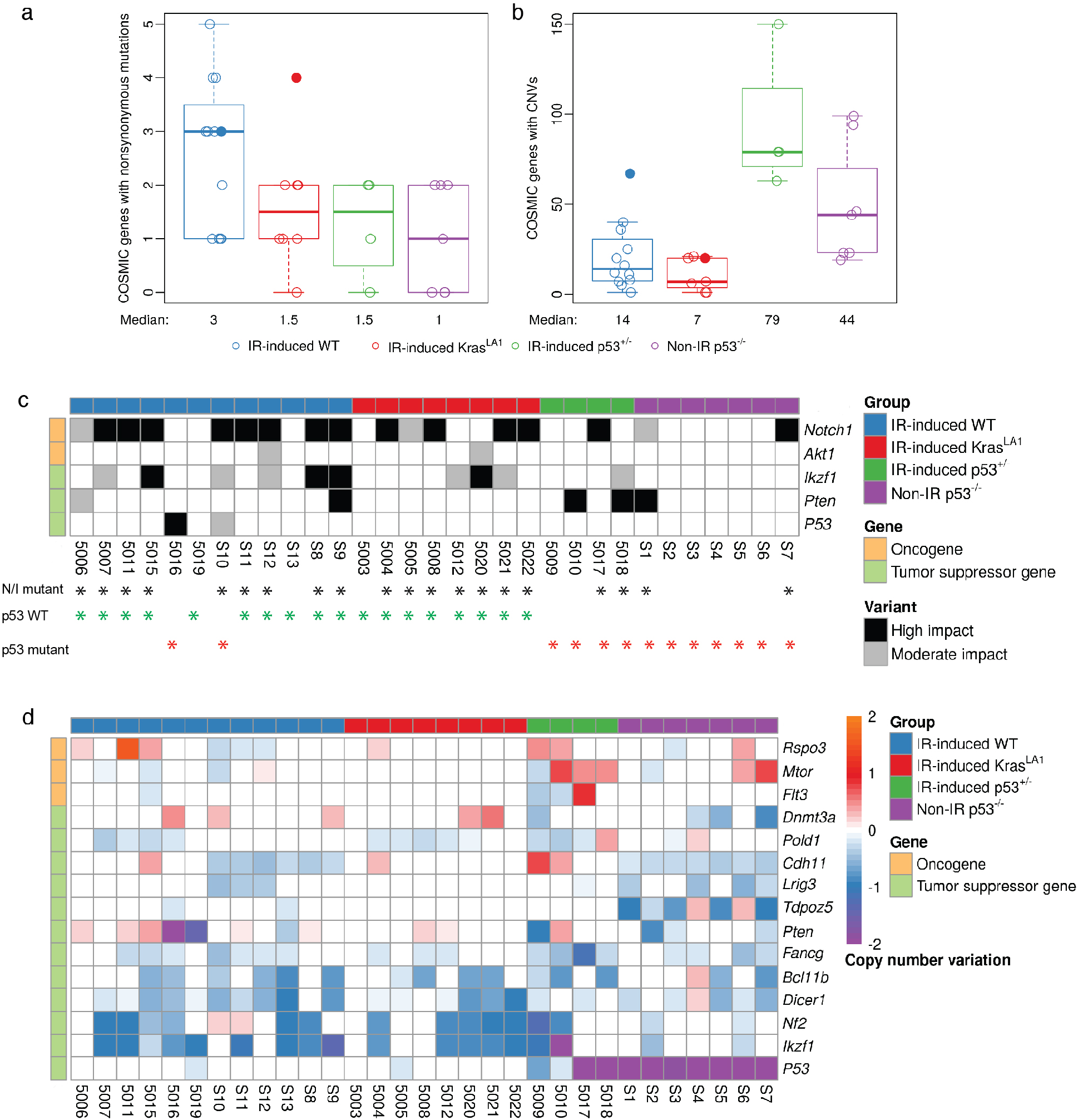
Somatic genetic alterations in COSMIC genes across murine thymic lymphomas. **a-b,** The number of COSMIC genes affected by nonsynonymous mutations and CNVs per tumor, respectively. Data from lymphomas that had substantially higher mutational loads (5015 and 5020) were denoted with closed circles. **c,** Recurrent mutations in COSMIC genes across thymic lymphomas. All recurrent mutations were validated by Sanger sequencing. Black asterisk: tumors with either *Notch1* and/or *Ikzf1* mutations (I/N mutant); Green asterisk: tumors with wildtype p53 (n=18); Red asterisk: tumors with *p53* mutations or harvested from p53^+/-^ or p53^-/-^ mice (n=13). **d,** COSMIC oncogenes that show a mean copy number gain and COSMIC tumor suppressor genes that show a mean copy number loss in thymic lymphomas. Genes are ordered within gene type by mean copy number across all samples. All panels illustrate the data for n=31 tumors.

We next examined individual COSMIC genes affected by nonsynonymous somatic mutations and/or CNVs across lymphomas from mice from the four genotypes (**Table S2**). Somatic mutations identified by WES were validated using Sanger Sequencing (**Table S3**). Our data revealed genotype-specific genetic alterations in signaling pathways, including Notch signaling, PI3K/AKT/mTOR signaling, MEK/ERK signaling, epigenetic modifiers, Hippo pathway, Wnt signaling and the DNA damage response (**Table S2**). Remarkably, among lymphomas that arose in irradiated WT and Kras^LA1^ mice, 90% (18/20) of them retained WT p53 (**Table S2 and Figure 3c**). Only two of these lymphomas harbored mutations in p53: tumors 5016 (missense mutation) and S10 (splicing mutation). Furthermore, we observed that p53 WT and p53 deficient lymphomas harbored distinct patterns of mutations that affect Notch signaling. In lymphomas that retained WT p53, approximately 83% (15/18) of these tumors harbored mutations in *Notch1* and/or *Ikzf1,* a negative regulator of Notch1 signaling (**Figure 3b and S4**). Three p53 WT lymphomas that did not harbor mutations in either *Notch1* or *Ikzf1*: tumors 5019 (mutations in *Ptpn11, Stil, Bcl11b* and *Ptprb),* S13 (mutation in *Ezh2)* and 5003 (Mutation in *Kras)* (**Figure S4**). In contrast to p53 WT lymphomas, mutations in *Notch1* or *Ikzf1* only occurred in around 38% (5/13) of p53 deficient lymphomas. (chi-squared test of proportions p-value=0.0281, **Figure 3c and S4**.) Nonsynonymous mutations in *Notch1* were exclusively found in exon 27 that encodes the heterodimerization domain or HD (all missense mutations) and exon 34 that encodes the proline, glutamic acid, serine, threonine-rich or PEST domain (missense, stop-gain and/or frameshift mutations). Also, nonsynonymous mutations in *Ikzf1* were exclusively found in exons 5/6 that encode the DNA binding domain (all missense mutations) and exon 9 (stop-gain or frameshift mutations) (**Figure S5**). These mutations recapitulate hotspot mutations in human *NOTCH1* and *IKZF1* observed in acute lymphoblastic leukemia patients^29,30^.

Similar to our findings from somatic mutational analysis of COSMIC genes, we observed a distinct difference in CNVs of *Ikzf1* between p53 WT and p53 deficient lymphomas (**Figure 3d**). While approximately 72% (13/18) of lymphomas that retained wild-type p53 show loss of *Ikzf1,* loss of *Ikzf1* was found in only 38% (5/13) of p53 deficient lymphomas. CNVs of *Ikzf1* were validated using a genomic qPCR assay (**Figure S6**). In contrast, *Mtor,* a critical gene that regulates cell growth and proliferation^31^, commonly showed copy number gains in tumors from mice with germline p53 deletion (5/11) (**Figure 3d**). Collectively, our analysis of somatic mutations and CNVs reveal distinct genetic alterations that occur in thymic lymphomas with and without wild-type p53. In particular, our findings demonstrate that p53 wild-type thymic lymphomas exhibit a high frequency of genetic alterations in *Notch1* and *Ikzf1,* two critical genes that regulate the Notch1 signaling pathway.

### Notch1 signaling is dysregulated primarily in thymic lymphomas harboring wild-type p53

Our findings of frequent *Notch1* pathway mutations in radiation-induced thymic lymphomas that retained wild-type p53 prompted us to further examine Notch1 signaling among the different lymphoma cohorts. One critical consequence of *Notch1* and *Ikzf1* mutations is increased expression of the intracellular domain of Notch1 (ICN)^32^. RT-PCR showed that mRNA of exon 32 - 33, which encodes for part of the ICN protein, was significantly overexpressed only in IR-induced WT and IR-induced Kras^LA1^ thymic lymphomas compared to normal thymuses from mice of the same genotype (Mann-Whitney U p-values 0.0080 and 0.0238, respectively, **Figure 4a**). Also, ICN protein was overexpressed in 100% of IR-induced WT and IR-induced Kras^LA1^ thymic lymphomas that we examined by Western blot (**Figure 4b and Figure S7**). In contrast, overexpression of ICN protein was only detected in 50% of IR-induced p53^+/-^ and 33% of Non-IR p53^-/-^ thymic lymphomas that we examined (**Figure 4b and Figure S7**). To determine the biological effects of ICN protein overexpression on the activation of Notch1 signaling, we quantified the mRNA expression of several transcriptional targets of *Notch1*. We observed that low-ICN expressing lymphomas, normal thymuses, and lymphomas expressing high ICN at the protein level not only exhibited differential mRNA expression of *3’ Notch1,* but also the Notch1 transcriptional targets *Hes1, Dtx1* and *Myc* (Kruskal-Wallace p-values 0.0002, 0.0335, 0.0009, 0.0002, respectively, **Figure 4c**).

**Figure 4.**
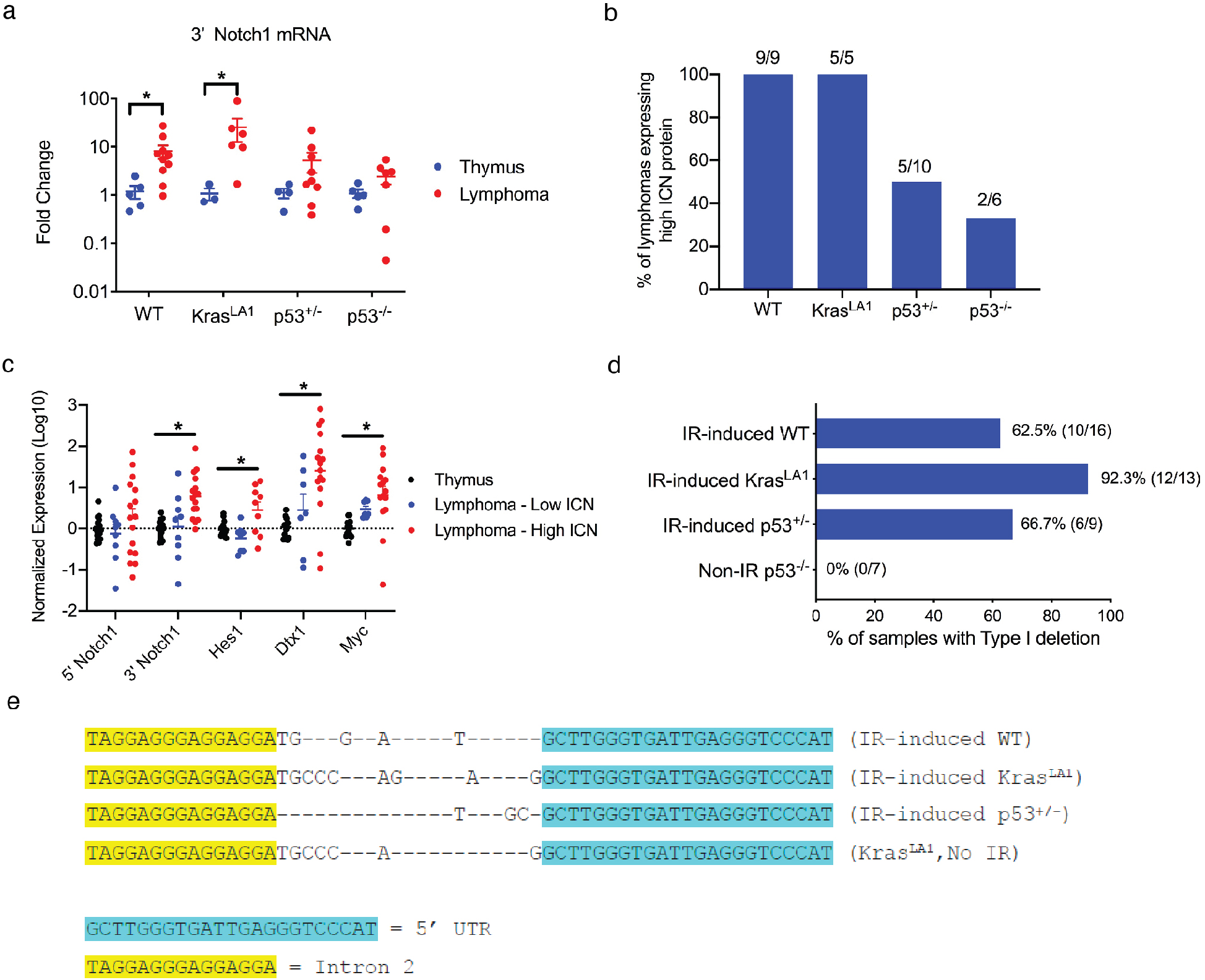
The Notch1 signaling pathway is upregulated in thymic lymphomas harboring functional p53. **a,** mRNA expression of intracellular domain of Notch1 (ICN). Results were compared between lymphomas and control thymuses from mice of the same genotype. Each dot represents one mouse. \**P*<0.05 by Mann-Whitney U test. **b,** The percentage of lymphomas expressing high ICN protein examined by Western blot. **c,** The mRNA expression of 5’ and 3’ regions of *Notch1* as well as transcriptional targets of Notch1. Each dot represents one mouse. \**P*<0.05 by Kruskal-Wallis test. **d,** Detection of Type 1 deletions of the *Notch1* gene among thymic lymphomas of different genotypes. **e,** Sequencing results of the type 1 deletion PCR product. Sanger sequencing showed that type 1 deletion resulted from a specific deletion recombination between the 5’ UTR and Intron 2. The sequences of *Notch1* type 1 deletion were conserved among thymic lymphomas across the genotypes of all three radiation-induced lymphoma models.

It has been shown that high ICN protein expression in mouse thymic lymphomas can be attributed to deletions within the *Notch1* locus. For example, small deletions of exons 1 and 2 of *Notch1,* termed Type I deletions, can result in overexpression of a truncated 3’ mRNA transcript from a cryptic promoter in exon 25 that is translated into the ICN protein^33^. To explore this potential mechanism underlying Notch1 activation, we performed PCR to detect Type I deletions of *Notch1* in our tumors (**Figure 4, d and e; Figure S8**). We observed that over 90% (12/13) of IR-induced Kras^LA1^ thymic lymphomas harbored Type I deletions, while Type I deletions were detected in around 60 to 70% of IR-induced WT lymphomas (10/16) and IR-induced p53^+/-^ lymphomas (6/9) (**Figure 4d**). Interestingly, Type I deletions were not detected in any of the Non-IR p53^-/-^ thymic lymphomas that we examined (0/7) (**Figure 4d**). The presence of type I deletions in p53 WT thymic lymphomas from unirradiated Kras^LA1^ mice indicates that the lack of Type I deletions in Non-IR p53^-/-^ lymphomas is unlikely to be a consequence of a lymphoma developing in the absence of radiation exposure. Instead, the lack of Type I deletions in Non-IR p53^-/-^ lymphomas is more likely a consequence of tumors forming in cells lacking p53 (**Figure 4e and Figure S8**).

Collectively, our results demonstrate that Notch1-mediated signaling is dysregulated at the DNA, RNA and protein levels in all IR-induced WT and IR-induced Kras^LA1^ thymic lymphomas that we examined. In contrast, only a subset of IR-induced p53^+/-^ and Non-IR p53^-/-^ thymic lymphomas that we studied showed aberrant activation of Notch1. These findings reveal that Notch1 signaling is dysregulated predominately in thymic lymphomas harboring WT p53.

### Inhibition of Notch1 signaling prevents the formation of radiation-induced thymic lymphoma in wild-type mice

To identify cells that express activated Notch1 signaling *in vivo,* we induced thymic lymphomas in Rbpj-Venus reporter mice^22^. Rbpj is a transcriptional co-activator of Notch1^34^, and cells positive for Venus in this model have been shown to exhibit elevated Notch1 activity^22^. In fully developed radiation-induced WT thymic lymphomas, we found that the majority of tumor cells expressed Venus (**Figure 5a**). Notably, Venus^+^ cells formed a distinct population that expressed surface markers CD4 and CD8 as well as CD44 (**Figure 5, b to c**). In parallel to the reporter assays, we examined thymocyte self-renewal using a thymocyte transplantation assay. We harvested thymocytes from WT mice 16 weeks after TBI and age-matched unirradiated controls and transplanted them into irradiated recipient mice (**Figure S9a**). The number of thymocytes in recipient mice 21 days after transplantation was significantly higher from irradiated donors (Mann-Whitney U p-value = 0.0238, **Figure S9b**). Moreover, flow cytometry analysis indicated that engraftment of donor thymocytes into the recipient thymus was much more successful from irradiated donors compared to thymocytes from unirradiated donors (Mann-Whitney U p-value = 0.0275, **Figure S9c**). Analysis of cell surface markers of thymocytes indicated that engrafted thymocytes were almost exclusively CD4^+^CD8^+^ and CD44^+^ (**Figure S9, d to f**). To determine the activity of Nocth1 signaling in self-renewing thymocytes, we performed thymocyte transplantation using thymocytes harvested from Rbpj-Venus reporter mice, backcrossed to a B6.SJL background, 19 weeks after TBI. We observed that the majority of donor-derived self-renewing thymocytes were Venus^+^ (**Figure 5d**). Further analysis of Venus^+^ self-renewing thymocytes showed a significant enrichment of CD44^+^ cells (**Figure 5, e and f**). Together, our findings from Rbpj-Venus reporter mice and *in vivo* thymocyte self-renewal assays indicate an important role of the Notch1-CD44 axis in regulating self-renewal of irradiated thymocytes *in vivo*.

**Figure 5.**
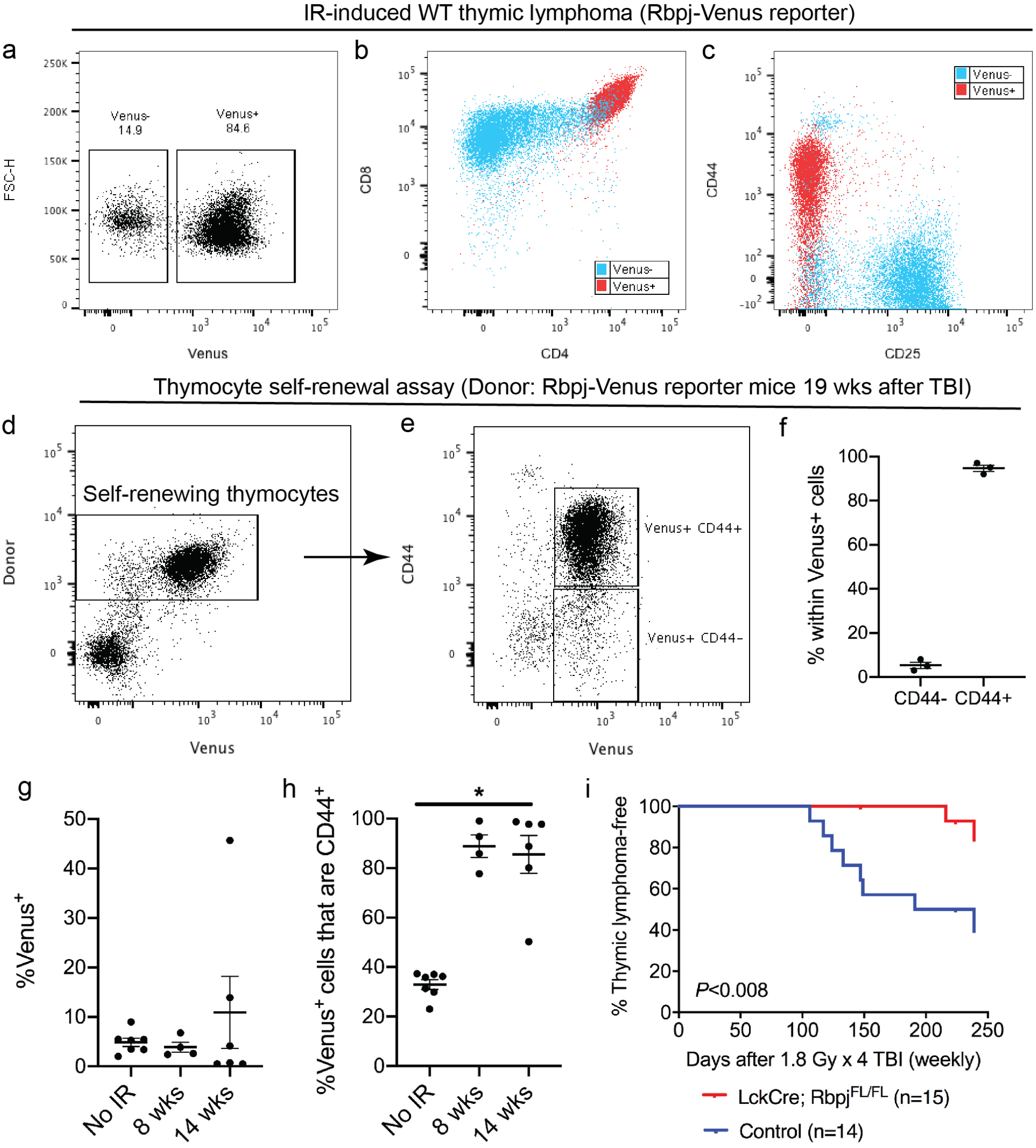
Notch1 signaling as the driver of the formation of radiation-induced thymic lymphoma in p53 WT mice. **a-c,** Representative flow cytometry data of radiation-induced thymic lymphoma that developed in Rbpj-Venus reporter mice on a p53 WT background. Venus^+^ cells and Venus^-^ cells were labeled in red and blue, respectively. **d-f,** Self-renewal of thymocytes harvested from Rbpj-Venus reporter mice 19 weeks post-irradiation. Donor thymocytes were injected intravenously into C57BL/6J recipients (CD45.1) 6 hours after 5 Gy TBI. Thymocytes were harvested from recipients 21 days after transplantation. Donor-derived Venus^+^ thymocytes were predominately CD44^+^. Data are presented as mean ± SE. Each dot represents one mouse. **g,** The percentage of Venus^+^ cells among total thymocytes harvested from Rbpj-Venus reporter mice at various time points after 1.8 Gy x 4 TBI. **h,** The percentage of Venus^+^ thymocytes also positive for CD44. Each dot presents one mouse. \**P*=0.0029 by Kruskal-Wallace test. **i,** Thymic lymphoma-free survival of LckCre; Rbpj^FL/FL^ mice and littermate controls that consist of Rbpj^FL/WT^ or Rbpj^FL/FL^ (No Cre) mice and LckCre; Rbpj^FL/WT^ mice that retain the expression of one Rbpj allele. All mice were exposed to 1.8 Gy x 4 TBI. *P*=0.0077 by log-rank test.

To examine the distribution of Venus^+^ cells at earlier stages of radiation-induced lymphomagenesis, we quantified the percentage of Venus^+^ thymocytes from unirradiated Rbpj-Venus reporter mice and Rbpj-Venus reporter mice 8 and 14 weeks after irradiation. Although the percentage of Venus^+^ thymocytes was not increased at 8 and 14 weeks after irradiation (**Figure 5g**), the percentage of Venus^+^ cells that are also CD44^+^ was significantly increased 8 and 14 weeks after irradiation (Kruskal-Wallace p-value = 0.0029, **Figure 5h**). These data suggest that while TBI did not cause an overall increase in Venus^+^ cells at earlier stages of lymphomagenesis, the subset of CD44^+^ thymocytes with activation of the Notch1 pathway as measured by Venus^+^ reporter expression was significantly enriched by irradiation.

To determine if Notch1-mediated signaling is necessary to drive the formation of radiation-induced thymic lymphoma, we performed genetic loss-of-function experiments to inhibit Notch1 signaling specifically in thymocytes. We generated LckCre; Rbpj floxed mice in which the Rbpj allele is specifically deleted in immature thymocytes beyond the DN2 stage^20,21^. LckCre; Rbpj^FL/FL^ mice and their littermate controls including Rbpj^FL/FL^ and Rbpj^FL/WT^ (No Cre) mice in addition to LckCre; Rbpj ^FL/WT^ mice that retain one Rbpj were exposed to 1.8 Gy x 4 TBI and followed for the formation of thymic lymphoma. Our results showed that the development of radiation-induced thymic lymphoma was significantly inhibited in LckCre; Rbpj^FL/FL^ mice (logrank p-value = 0.0077, **Figure 5i**). Together, our findings from genetic analyses of lymphomas, Rbpj-Venus reporter mice experiments, thymocyte self-renewal assays and thymocyte-specific Rbpj knockout mice demonstrate that activated Notch1 signaling drives the development of radiation-induced thymic lymphoma in p53 wild-type mice.

## DISCUSSION

In the present study, we conducted a comprehensive analysis of somatic mutations and CNVs in thymic lymphomas that developed in irradiated wild-type, Kras^LA1^, p53^+/-^ mice as well as in unirradiated mice lacking both alleles of p53 in hematopoietic cells. Our results reveal that murine radiation-induced thymic lymphomas harbor recurrent mutations in putative driver genes that have been reported in human T-cell acute lymphoblastic leukemia (T-ALL), including *Pten, Akt1, Ikzf1* and *Notch1*^29^ (**Figure 3**). In particular, we found that mutations in the *Notch1* oncogene frequently occur in radiation-induced thymic lymphomas harboring wild-type p53. Although the presence of mutations and CNVs that affect the Notch1 signaling pathway in murine radiation-induced thymic lymphomas have been reported previously^35–37^, our results from mice lacking functional *Rbpj* in immature thymocytes, for the first time, demonstrate that activation of Notch1 signaling is necessary to drive radiation-induced thymic lymphomas in p53 wild-type mice (**Figure 5**). Our study is distinct from papers that investigate the role of Notch1 in T-cell leukemogenesis by overexpressing ICN because the development of thymic lymphomas in our model is driven by genetic alterations that spontaneously occur in wild-type mice following TBI. Together, our findings uncover the mutational landscape of murine radiation-induced thymic lymphomas and define the pivotal role of activated Notch1 signaling in driving tumor development in p53 wild-type mice.

Our study also provides novel insights into how TBI selects for Notch1-mutant thymic lymphomas in p53 wild-type mice. Our results from Rbpj-Venus reporter mice indicate that malignant thymocytes expressing high Notch1 activity form a distinct population positive for CD44. In addition, we observed that irradiated thymocytes that undergo self-renewal *in vivo* are almost exclusively positive for CD44^+^ (**Figure 5**). Intriguingly, the results from our time-course study show that TBI does not cause an overall increase in Venus^+^ cells at 8 and 14 weeks after irradiation, however, the subset of CD44^+^ thymocytes with activation of the Notch1 pathway as measured by Venus^+^ reporter expression was significantly enriched by TBI at both time points. The expansion of CD44^+^ thymocytes after TBI was also observed in an independent study^38^. Moreover, inhibition of CD44 significantly suppresses the engraftment of Notch1-mutant human T-ALL cells^39^. Together, these results suggest that the Notch1-CD44 axis plays an important role in regulating self-renewal and subsequent malignant transformation of irradiated thymocytes *in vivo*.

Our results also demonstrate that alterations in Notch1-mediated signaling occur more frequently in p53 wild-type thymic lymphomas following TBI than in lymphomas that develop in irradiated p53^+/-^ or Non-IR p53^-/-^ mice. Indeed, while 100% of thymic lymphomas that develop in p53 wild-type mice following TBI overexpress ICN, overexpression of ICN only occurs in 50% of radiation-induced thymic lymphomas that develop in p53^+/-^ mice (**Figure 4**). These results are consistent with the findings that *Notch1* PEST mutations are detected in approximately 40% of thymic lymphomas that develop in p53^+/-^ mice following a single dose of 4 Gy TBI^40^. The frequency of ICN overexpression is even lower in radiation naïve thymic lymphomas driven by the loss of p53 (33%) (**Figure 4**). This inverse relationship between mutations in *Notch1* and *p53* has also been reported in murine thymic lymphomas generated by viral insertion^41^. In this paper, Uren and colleagues found that mutations that cause overexpression of ICN are almost exclusively detected in tumors harboring wild-type p53 and p19^Arf^. One possible explanation for this phenomenon is that p53 deficient mice have impaired expression Fbxw7, a transcriptional target of p53 that regulates the turnover of ICN^42,43^. Indeed, *Notch1* PEST domain mutations are not detected in radiation-induced thymic lymphomas of p53^+/-^; Fbxw7^+/-^ mice^40^. However, as Notch1 and p53 both regulate the expression of numerous transcriptional targets, further investigation is warranted to dissect how Notch1 and p53 signaling pathways interact to regulate the formation of T-cell lymphoblastic lymphoma/leukemia.

Our mutational analyses using whole-exome sequencing data found no statistical evidence of a difference in the number of nonsynonymous somatic mutations per tumor between radiation-induced thymic lymphomas and thymic lymphomas that develop in p53^-/-^ mice without irradiation (**Figure 1**). Similarly, analyses of mutational signatures by assessing the proportion of indels, the distribution of SNVs and cosine-similarity with COSMIC mutational signatures of human cancers do not show a significant difference between radiation-induced thymic lymphomas and Non-IR p53^-/-^ thymic lymphomas (**Figure 1 and S3**). While the absence of evidence for these differences may merely be due to the small size of the experimental populations, a study by Wang et al. also reported that therapy-related myeloid neoplasms (t-MNs) and de novo myeloid neoplasms in human patients exhibit a similar number of somatic mutations and single nucleotide substitutions^44^. In contrast to blood cancers, solid tumors that developed in humans and mice following focal irradiation display unique mutational processes compared to radiation naïve solid tumors. For example, radiation-induced sarcomas exhibit a strong oxidative mutation signature as shown by a preference of C to T and the reverse (G to A) single nucleotide substitutions as well as harboring a higher proportion of nonsynonymous deletion^10^. However, all of these unique mutational signatures were not observed in radiation-induced thymic lymphomas. Besides somatic mutations, we found that the number of genes affected by CNVs is markedly higher in thymic lymphomas in p53^+/-^ mice treated with four weekly fractions of 1.8 Gy TBI (**Figure 2**). A similar increase in genomic instability was also reported in thymic lymphomas of p53^+/-^ mice after a single fraction of 4 Gy TBI^45^. Collectively, these genomic studies provide compelling evidence to support a model in humans and mice without germline mutations where the formation of hematologic malignancies following exposure to ionizing radiation occurs through a non-cell-autonomous mechanism^8,46,47^. However, for individuals harboring germline mutations in DNA damage response genes, such as *p53* and *Mlh1*, ionizing radiation may cause blood cancers through a more direct cell-autonomous effect by increasing genomic instability in tumor-initiating cells^13,14,45^.

In sum, the results in the present study reveal the mutational landscape of coding genes and identify key drivers in murine radiation-induced thymic lymphoma, a classic model that has been used to study radiation carcinogenesis for over 70 years. These findings lay the foundation for a better understanding of mechanisms underlying multi-step carcinogenesis of hematologic malignances following total-body irradiation.

## ACKNOWLEDGMENT

This work was supported by the following grants: National Cancer Institute (R35CA197616 to DGK, K99CA212198 and R00CA212198 to C-LL), Whitehead Scholar Award from Duke University School of Medicine (C-LL), and the Duke Cancer Center Support Grant 5P30CA14236-44.

## AUTHOR CONTRIBUTIONS

C-LL and DGK designed the study. C-LL, KDB, SH, IC, LL, and ARD performed *in vitro* and *in vivo* mouse experiments. DZ, ABS, JG and XQ performed bioinformatic analyses. C-LL, KDB, DZ, ABS, XQ, IC, ARD, MJH, KO, and DGK analyzed and interpreted data. C-LL, ARD, and DGK wrote the manuscript. All authors edited and approved the manuscript.

## REFERENCES

1 Smith, S. M., Le Beau, M. M., Huo, D., Karrison, T., Sobecks, R. M., Anastasi, J., Vardiman, J. W., Rowley, J. D. & Larson, R. A. Clinical-cytogenetic associations in 306 patients with therapy-related myelodysplasia and myeloid leukemia: the University of Chicago series. Blood 102, 43–52, doi:10.1182/blood-2002-11-3343 (2003).

2 Shuryak, I., Sachs, R. K., Hlatky, L., Little, M. P., Hahnfeldt, P. & Brenner, D. J. Radiation-induced leukemia at doses relevant to radiation therapy: modeling mechanisms and estimating risks. J Natl Cancer Inst 98, 1794–1806, doi:10.1093/jnci/djj497 (2006).

3 Ozasa, K., Shimizu, Y., Suyama, A., Kasagi, F., Soda, M., Grant, E. J., Sakata, R., Sugiyama, H. & Kodama, K. Studies of the mortality of atomic bomb survivors, Report 14, 1950-2003: an overview of cancer and noncancer diseases. Radiat Res 177, 229–243 (2012).

4 Hsu, W. L., Preston, D. L., Soda, M., Sugiyama, H., Funamoto, S., Kodama, K., Kimura, A., Kamada, N., Dohy, H., Tomonaga, M., Iwanaga, M., Miyazaki, Y., Cullings, H. M., Suyama, A., Ozasa, K., Shore, R. E. & Mabuchi, K. The incidence of leukemia, lymphoma and multiple myeloma among atomic bomb survivors: 1950–2001. Radiat Res 179, 361–382, doi:10.1667/RR2892.1 (2013).

5 Rivina, L. & Schiestl, R. Mouse models of radiation-induced cancers. Advances in genetics 84, 83–122, doi:10.1016/B978-0-12-407703-4.00003-7 (2013).

6 Kaplan, H. S. Observations on radiation-induced lymphoid tumors of mice. Cancer Res 7, 141–147 (1947).

7 Kaplan, H. S. & Brown, M. B. A quantitative dose-response study of lymphoid-tumor development in irradiated C 57 black mice. J Natl Cancer Inst 13, 185–208 (1952).

8 Lee, C. L., Castle, K. D., Moding, E. J., Blum, J. M., Williams, N., Luo, L., Ma, Y., Borst, L. B., Kim, Y. & Kirsch, D. G. Acute DNA damage activates the tumour suppressor p53 to promote radiation-induced lymphoma. Nat Commun 6, 8477, doi:10.1038/ncomms9477 (2015).

9 Lee, C. L., Mowery, Y. M., Daniel, A. R., Zhang, D., Sibley, A. B., Delaney, J. R., Wisdom, A. J., Qin, X., Wang, X., Caraballo, I., Gresham, J., Luo, L., Van Mater, D., Owzar, K. & Kirsch, D. G. Mutational landscape in genetically engineered, carcinogen-induced, and radiation-induced mouse sarcoma. JCI Insight 4, doi:10.1172/jci.insight.128698 (2019).

10 Sherborne, A. L., Davidson, P. R., Yu, K., Nakamura, A. O., Rashid, M. & Nakamura, J. L. Mutational Analysis of Ionizing Radiation Induced Neoplasms. Cell Rep 12, 1915–1926, doi:10.1016/j.celrep.2015.08.015 (2015).

11 Thibodeau, B. J., Lavergne, V., Dekhne, N., Benitez, P., Amin, M., Ahmed, S., Nakamura, J. L., Davidson, P. R., Nakamura, A. O., Grills, I. S., Chen, P. Y., Wobb, J. & Wilson, G. D. Mutational landscape of radiation-associated angiosarcoma of the breast. Oncotarget 9, 10042–10053, doi:10.18632/oncotarget.24273 (2018).

12 Behjati, S., Gundem, G., Wedge, D. C., Roberts, N. D., Tarpey, P. S., Cooke, S. L., Van Loo, P., Alexandrov, L. B., Ramakrishna, M., Davies, H., Nik-Zainal, S., Hardy, C., Latimer, C., Raine, K. M., Stebbings, L., Menzies, A., Jones, D., Shepherd, R., Butler, A. P., Teague, J. W., Jorgensen, M., Khatri, B., Pillay, N., Shlien, A., Futreal, P. A., Badie, C., Group, I. P., McDermott, U., Bova, G. S., Richardson, A. L., Flanagan, A. M., Stratton, M. R. & Campbell, P. J. Mutational signatures of ionizing radiation in second malignancies. Nat Commun 7, 12605, doi:10.1038/ncomms12605 (2016).

13 Patel, R., Zhang, L., Desai, A., Hoenerhoff, M. J., Kennedy, L. H., Radivoyevitch, T., Ban, Y., Chen, X. S., Gerson, S. L. & Welford, S. M. Mlh1 deficiency increases the risk of hematopoietic malignancy after simulated space radiation exposure. Leukemia 33, 1135–1147, doi:10.1038/s41375-018-0269-8 (2019).

14 Rose Li, Y., Halliwill, K. D., Adams, C. J., Iyer, V., Riva, L., Mamunur, R., Jen, K. Y., Del Rosario, R., Fredlund, E., Hirst, G., Alexandrov, L. B., Adams, D. & Balmain, A. Mutational signatures in tumours induced by high and low energy radiation in Trp53 deficient mice. Nat Commun 11, 394, doi:10.1038/s41467-019-14261-4 (2020).

15 Johnson, L., Mercer, K., Greenbaum, D., Bronson, R. T., Crowley, D., Tuveson, D. A. & Jacks, T. Somatic activation of the K-ras oncogene causes early onset lung cancer in mice. Nature 410, 1111–1116, doi:10.1038/3507412935074129 [pii] (2001).

16 Jacks, T., Remington, L., Williams, B. O., Schmitt, E. M., Halachmi, S., Bronson, R. T. & Weinberg, R. A. Tumor spectrum analysis in p53-mutant mice. Curr Biol 4, 1–7 (1994).

17 Kemp, C. J., Wheldon, T. & Balmain, A. p53-deficient mice are extremely susceptible to radiation-induced tumorigenesis. Nat Genet 8, 66–69, doi:10.1038/ng0994-66 (1994).

18 Moding, E. J., Min, H. D., Castle, K. D., Ali, M., Woodlief, L., Williams, N., Ma, Y., Kim, Y., Lee, C. L. & Kirsch, D. G. An extra copy of p53 suppresses development of spontaneous Kras-driven but not radiation-induced cancer. JCI Insight 1, doi:10.1172/jci.insight.86698 (2016).

19 Lee, C. L., Moding, E. J., Cuneo, K. C., Li, Y., Sullivan, J. M., Mao, L., Washington, I., Jeffords, L. B., Rodrigues, R. C., Ma, Y., Das, S., Kontos, C. D., Kim, Y., Rockman, H. A. & Kirsch, D. G. p53 functions in endothelial cells to prevent radiation-induced myocardial injury in mice. Sci Signal 5, ra52, doi:10.1126/scisignal.2002918 (2012).

20 Hennet, T., Hagen, F. K., Tabak, L. A. & Marth, J. D. T-cell-specific deletion of a polypeptide N-acetylgalactosaminyl-transferase gene by site-directed recombination. Proc Natl Acad Sci U S A 92, 12070–12074, doi:10.1073/pnas.92.26.12070 (1995).

21 Tanigaki, K., Han, H., Yamamoto, N., Tashiro, K., Ikegawa, M., Kuroda, K., Suzuki, A., Nakano, T. & Honjo, T. Notch-RBP-J signaling is involved in cell fate determination of marginal zone B cells. Nat Immunol 3, 443–450, doi:10.1038/ni793 (2002).

22 Nowotschin, S., Xenopoulos, P., Schrode, N. & Hadjantonakis, A. K. A bright single-cell resolution live imaging reporter of Notch signaling in the mouse. BMC Dev Biol 13, 15, doi:10.1186/1471-213X-13-15 (2013).

23 Gerby, B., Tremblay, C. S., Tremblay, M., Rojas-Sutterlin, S., Herblot, S., Hebert, J., Sauvageau, G., Lemieux, S., Lecuyer, E., Veiga, D. F. & Hoang, T. SCL, LMO1 and Notch1 reprogram thymocytes into self-renewing cells. PLoS genetics 10, e1004768, doi:10.1371/journal.pgen.1004768 (2014).

24 R Core Team. R: A language and environment for statistical computing (Vienna, Austria, 2020).

25 Huber, W., Carey, V. J., Gentleman, R., Anders, S., Carlson, M., Carvalho, B. S., Bravo, H. C., Davis, S., Gatto, L., Girke, T., Gottardo, R., Hahne, F., Hansen, K. D., Irizarry, R. A., Lawrence, M., Love, M. I., MacDonald, J., Obenchain, V., Oles, A. K., Pages, H., Reyes, A., Shannon, P., Smyth, G. K., Tenenbaum, D., Waldron, L. & Morgan, M. Orchestrating high-throughput genomic analysis with Bioconductor. Nat Methods 12, 115–121, doi:10.1038/nmeth.3252 (2015).

26 Xie, Y. knitr: A Comprehensive Tool for Reproducible Research in R. Implement Reprod Res 1, 20 (2014).

27 Alexandrov, L. B., Nik-Zainal, S., Wedge, D. C., Aparicio, S. A., Behjati, S., Biankin, A. V., Bignell, G. R., Bolli, N., Borg, A., Borresen-Dale, A. L., Boyault, S., Burkhardt, B., Butler, A. P., Caldas, C., Davies, H. R., Desmedt, C., Eils, R., Eyfjord, J. E., Foekens, J. A., Greaves, M., Hosoda, F., Hutter, B., Ilicic, T., Imbeaud, S., Imielinski, M., Jager, N., Jones, D. T., Jones, D., Knappskog, S., Kool, M., Lakhani, S. R., Lopez-Otin, C., Martin, S., Munshi, N. C., Nakamura, H., Northcott, P. A., Pajic, M., Papaemmanuil, E., Paradiso, A., Pearson, J. V., Puente, X. S., Raine, K., Ramakrishna, M., Richardson, A. L., Richter, J., Rosenstiel, P., Schlesner, M., Schumacher, T. N., Span, P. N., Teague, J. W., Totoki, Y., Tutt, A. N., Valdes-Mas, R., van Buuren, M. M., van’t Veer, L., Vincent-Salomon, A., Waddell, N., Yates, L. R., Australian Pancreatic Cancer Genome, I., Consortium, I. B. C., Consortium, I. M.-S., PedBrain, I., Zucman-Rossi, J., Futreal, P. A., McDermott, U., Lichter, P., Meyerson, M., Grimmond, S. M., Siebert, R., Campo, E., Shibata, T., Pfister, S. M., Campbell, P. J. & Stratton, M. R. Signatures of mutational processes in human cancer. Nature 500, 415–421, doi:10.1038/nature12477 (2013).

28 Jiang, Y., Wang, R., Urrutia, E., Anastopoulos, I. N., Nathanson, K. L. & Zhang, N. R. CODEX2: full-spectrum copy number variation detection by high-throughput DNA sequencing. Genome Biol 19, 202, doi:10.1186/s13059-018-1578-y (2018).

29 Liu, Y., Easton, J., Shao, Y., Maciaszek, J., Wang, Z., Wilkinson, M. R., McCastlain, K., Edmonson, M., Pounds, S. B., Shi, L., Zhou, X., Ma, X., Sioson, E., Li, Y., Rusch, M., Gupta, P., Pei, D., Cheng, C., Smith, M. A., Auvil, J. G., Gerhard, D. S., Relling, M. V., Winick, N. J., Carroll, A. J., Heerema, N. A., Raetz, E., Devidas, M., Willman, C. L., Harvey, R. C., Carroll, W. L., Dunsmore, K. P., Winter, S. S., Wood, B. L., Sorrentino, B. P., Downing, J. R., Loh, M. L., Hunger, S. P., Zhang, J. & Mullighan, C. G. The genomic landscape of pediatric and young adult T-lineage acute lymphoblastic leukemia. Nat Genet 49, 1211–1218, doi:10.1038/ng.3909 (2017).

30 Lana, T., de Lorenzo, P., Bresolin, S., Bronzini, I., den Boer, M. L., Cave, H., Fronkova, E., Stanulla, M., Zaliova, M., Harrison, C. J., de Groot, H., Valsecchi, M. G., Biondi, A., Basso, G., Cazzaniga, G. & te Kronnie, G. Refinement of IKZF1 status in pediatric Philadelphia-positive acute lymphoblastic leukemia. Leukemia 29, 2107–2110, doi:10.1038/leu.2015.78 (2015).

31 Guertin, D. A. & Sabatini, D. M. Defining the role of mTOR in cancer. Cancer Cell 12, 9–22, doi:10.1016/j.ccr.2007.05.008 (2007).

32 Witkowski, M. T., Cimmino, L., Hu, Y., Trimarchi, T., Tagoh, H., McKenzie, M. D., Best, S. A., Tuohey, L., Willson, T. A., Nutt, S. L., Busslinger, M., Aifantis, I., Smyth, G. K. & Dickins, R. A. Activated Notch counteracts Ikaros tumor suppression in mouse and human T-cell acute lymphoblastic leukemia. Leukemia 29, 1301–1311, doi:10.1038/leu.2015.27 (2015).

33 Ashworth, T. D., Pear, W. S., Chiang, M. Y., Blacklow, S. C., Mastio, J., Xu, L., Kelliher, M., Kastner, P., Chan, S. & Aster, J. C. Deletion-based mechanisms of Notch1 activation in T-ALL: key roles for RAG recombinase and a conserved internal translational start site in Notch1. Blood 116, 5455–5464, doi:10.1182/blood-2010-05-286328 (2010).

34 Castel, D., Mourikis, P., Bartels, S. J., Brinkman, A. B., Tajbakhsh, S. & Stunnenberg, H. G. Dynamic binding of RBPJ is determined by Notch signaling status. Genes Dev 27, 1059–1071, doi:10.1101/gad.211912.112 (2013).

35 Tsuji, H., Ishii-Ohba, H., Ukai, H., Katsube, T. & Ogiu, T. Radiation-induced deletions in the 5’ end region of Notch1 lead to the formation of truncated proteins and are involved in the development of mouse thymic lymphomas. Carcinogenesis 24, 1257–1268, doi:10.1093/carcin/bgg071 (2003).

36 Lopez-Nieva, P., Santos, J. & Fernandez-Piqueras, J. Defective expression of Notch1 and Notch2 in connection to alterations of c-Myc and Ikaros in gamma-radiation-induced mouse thymic lymphomas. Carcinogenesis 25, 1299–1304, doi:10.1093/carcin/bgh124 (2004).

37 Ohi, H., Mishima, Y., Kamimura, K., Maruyama, M., Sasai, K. & Kominami, R. Multi-step lymphomagenesis deduced from DNA changes in thymic lymphomas and atrophic thymuses at various times after gamma-irradiation. Oncogene 26, 5280–5289, doi:10.1038/sj.onc.1210325 (2007).

38 Sen-Majumdar, A., Guidos, C., Kina, T., Lieberman, M. & Weissman, I. L. Characterization of preneoplastic thymocytes and of their neoplastic progression in irradiated C57BL/Ka mice. J Immunol 153, 1581–1592 (1994).

39 Garcia-Peydro, M., Fuentes, P., Mosquera, M., Garcia-Leon, M. J., Alcain, J., Rodriguez, A., Garcia de Miguel, P., Menendez, P., Weijer, K., Spits, H., Scadden, D. T., Cuesta-Mateos, C., Munoz-Calleja, C., Sanchez-Madrid, F. & Toribio, M. L. The NOTCH1/CD44 axis drives pathogenesis in a T cell acute lymphoblastic leukemia model. J Clin Invest 128, 2802–2818, doi:10.1172/JCI92981 (2018).

40 Jen, K. Y., Song, I. Y., Banta, K. L., Wu, D., Mao, J. H. & Balmain, A. Sequential mutations in Notch1, Fbxw7, and Tp53 in radiation-induced mouse thymic lymphomas. Blood, doi:10.1182/blood-2011-01-327619 (2011).

41 Uren, A. G., Kool, J., Matentzoglu, K., de Ridder, J., Mattison, J., van Uitert, M., Lagcher, W., Sie, D., Tanger, E., Cox, T., Reinders, M., Hubbard, T. J., Rogers, J., Jonkers, J., Wessels, L., Adams, D. J., van Lohuizen, M. & Berns, A. Large-scale mutagenesis in p19(ARF)- and p53-deficient mice identifies cancer genes and their collaborative networks. Cell 133, 727–741, doi:10.1016/j.cell.2008.03.021 (2008).

42 Mao, J. H., Perez-Losada, J., Wu, D., Delrosario, R., Tsunematsu, R., Nakayama, K. I., Brown, K., Bryson, S. & Balmain, A. Fbxw7/Cdc4 is a p53-dependent, haploinsufficient tumour suppressor gene. Nature 432, 775–779, doi:10.1038/nature03155 (2004).

43 Matsuoka, S., Oike, Y., Onoyama, I., Iwama, A., Arai, F., Takubo, K., Mashimo, Y., Oguro, H., Nitta, E., Ito, K., Miyamoto, K., Yoshiwara, H., Hosokawa, K., Nakamura, Y., Gomei, Y., Iwasaki, H., Hayashi, Y., Matsuzaki, Y., Nakayama, K., Ikeda, Y., Hata, A., Chiba, S., Nakayama, K. I. & Suda, T. Fbxw7 acts as a critical fail-safe against premature loss of hematopoietic stem cells and development of T-ALL. Genes & Development 22, 986–991, doi:10.1101/gad.1621808 (2008).

44 Wong, T. N., Ramsingh, G., Young, A. L., Miller, C. A., Touma, W., Welch, J. S., Lamprecht, T. L., Shen, D., Hundal, J., Fulton, R. S., Heath, S., Baty, J. D., Klco, J. M., Ding, L., Mardis, E. R., Westervelt, P., DiPersio, J. F., Walter, M. J., Graubert, T. A., Ley, T. J., Druley, T. E., Link, D. C. & Wilson, R. K. Role of TP53 mutations in the origin and evolution of therapy-related acute myeloid leukaemia. Nature 518, 552–555, doi:10.1038/nature13968 (2015).

45 Mao, J. H., Wu, D., Perez-Losada, J., Jiang, T., Li, Q., Neve, R. M., Gray, J. W., Cai, W. W. & Balmain, A. Crosstalk between Aurora-A and p53: frequent deletion or downregulation of Aurora-A in tumors from p53 null mice. Cancer Cell 11, 161–173, doi:10.1016/j.ccr.2006.11.025 (2007).

46 Marusyk, A., Casas-Selves, M., Henry, C. J., Zaberezhnyy, V., Klawitter, J., Christians, U. & DeGregori, J. Irradiation alters selection for oncogenic mutations in hematopoietic progenitors. Cancer research 69, 7262–7269, doi:10.1158/0008-5472.CAN-09-0604 (2009).

47 Fleenor, C. J., Marusyk, A. & DeGregori, J. Ionizing radiation and hematopoietic malignancies: altering the adaptive landscape. Cell Cycle 9, 3005–3011 (2010).

