## Supplementary Figures and Tables for "Activation of Notch1 drives the development of radiation-induced thymic lymphoma in p53 wild-type mice"

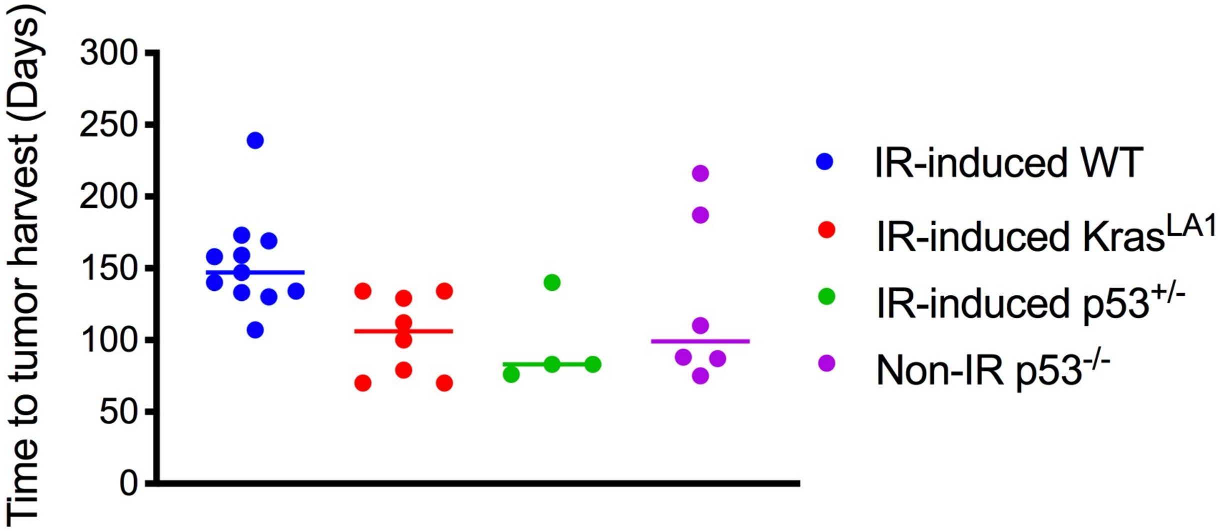


**Figure S1.** The latency period of thymic lymphomas from irradiated and unirradiated mice. Radiation-induced thymic lymphomas were generated in wild-type (WT) (C57BL/6J) mice, Kras$^{LA1}$ mice or p53$^{+/-}$ mice that were exposed to 1.8 Gy total-body irradiation (TBI) per week for 4 consecutive weeks. Unirradiated Tie2Cre; p53$^{FL/-}$ mice (Non-IR p53$^{-/-}$) in which both alleles of the p53 gene are deleted in endothelial cells and hematopoietic cells were used to generate thymic lymphomas in the absence of irradiation. For IR-induced WT, IR-induced Kras$^{LA1}$ and IR-induced p53$^{+/-}$ mice, data represent the number of days after exposure to 1.8 Gy x 4 TBI. For Non-IR p53$^{-/-}$ mice, data represent the number of days after the mice reached 7 weeks old. Data are presented as dot plots with median. Each dot represents one tumor.

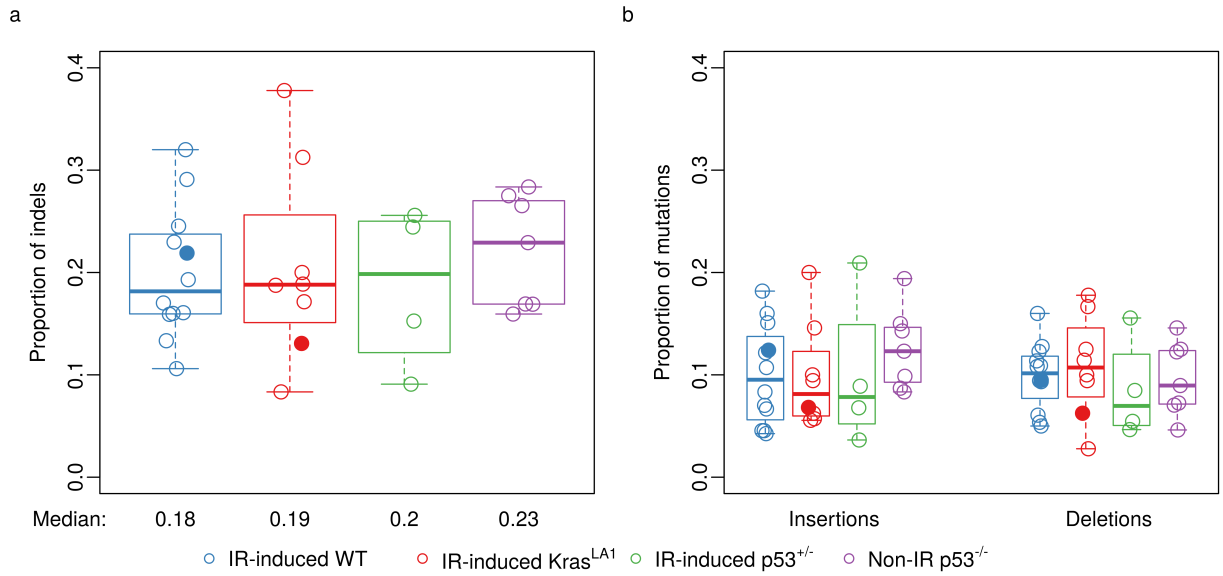


**Figure S2.** The distribution of sequence variants within total (synonymous and nonsynonymous) somatic mutations. **a,** The proportion of insertion-deletions (indels) within total mutations. **b,** The proportions of insertions and deletions within total mutations. Data from lymphomas that had a substantially higher number of somatic mutations (5015 and 5020) were denoted by closed circles.

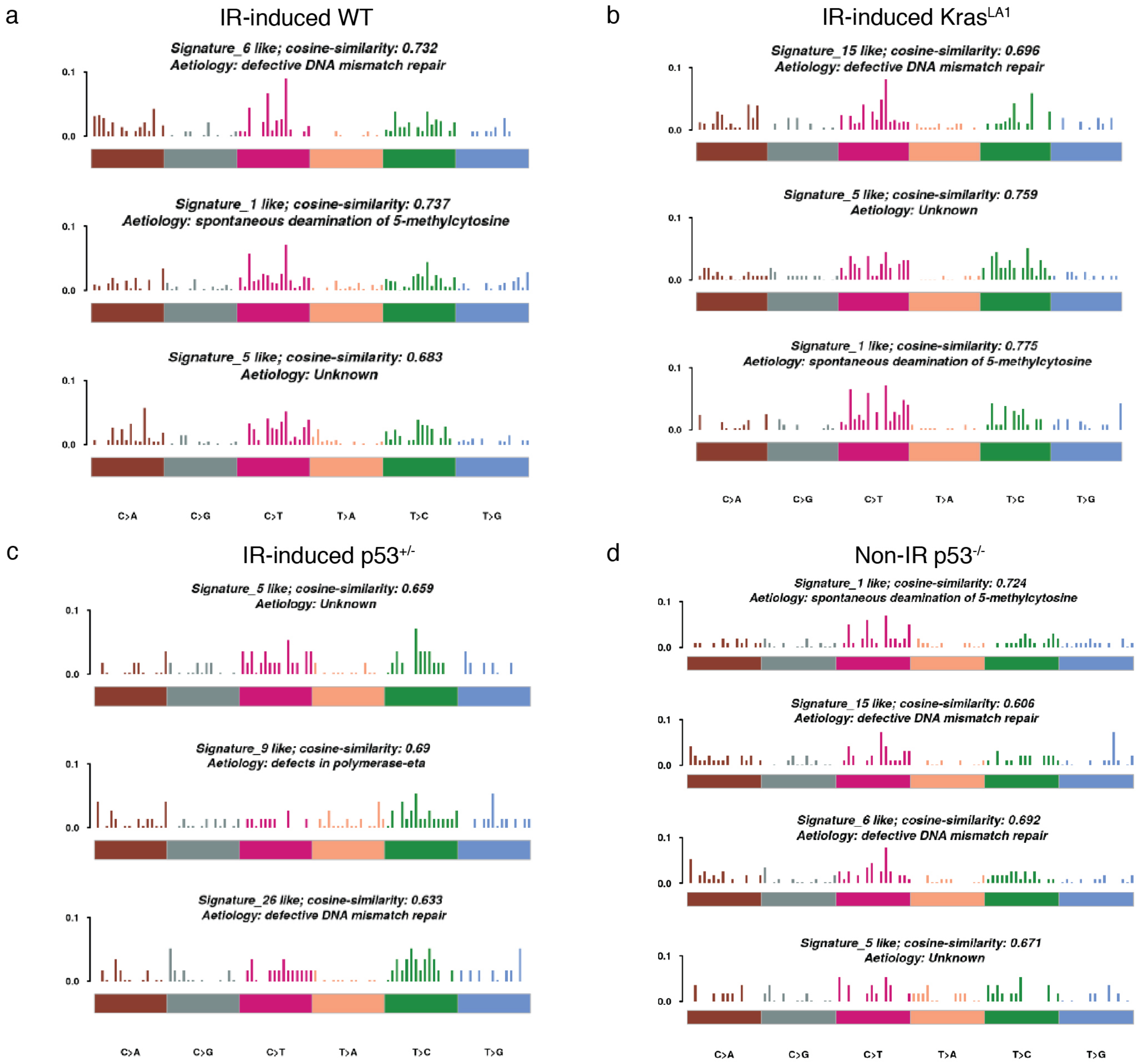


**Figure S3.** Mutational signature analysis of thymic lymphomas. Mutational signatures of each lymphoma genotype were generated using nonnegative matrix factorization (NMF) trinucleotide-based analysis. From each lymphoma genotype, either 3 or 4 individual signatures were generated. Each murine lymphoma signature was compared to COSMIC mutational signatures of human cancers to identify the COSMIC signature with the highest cosine-similarity score.

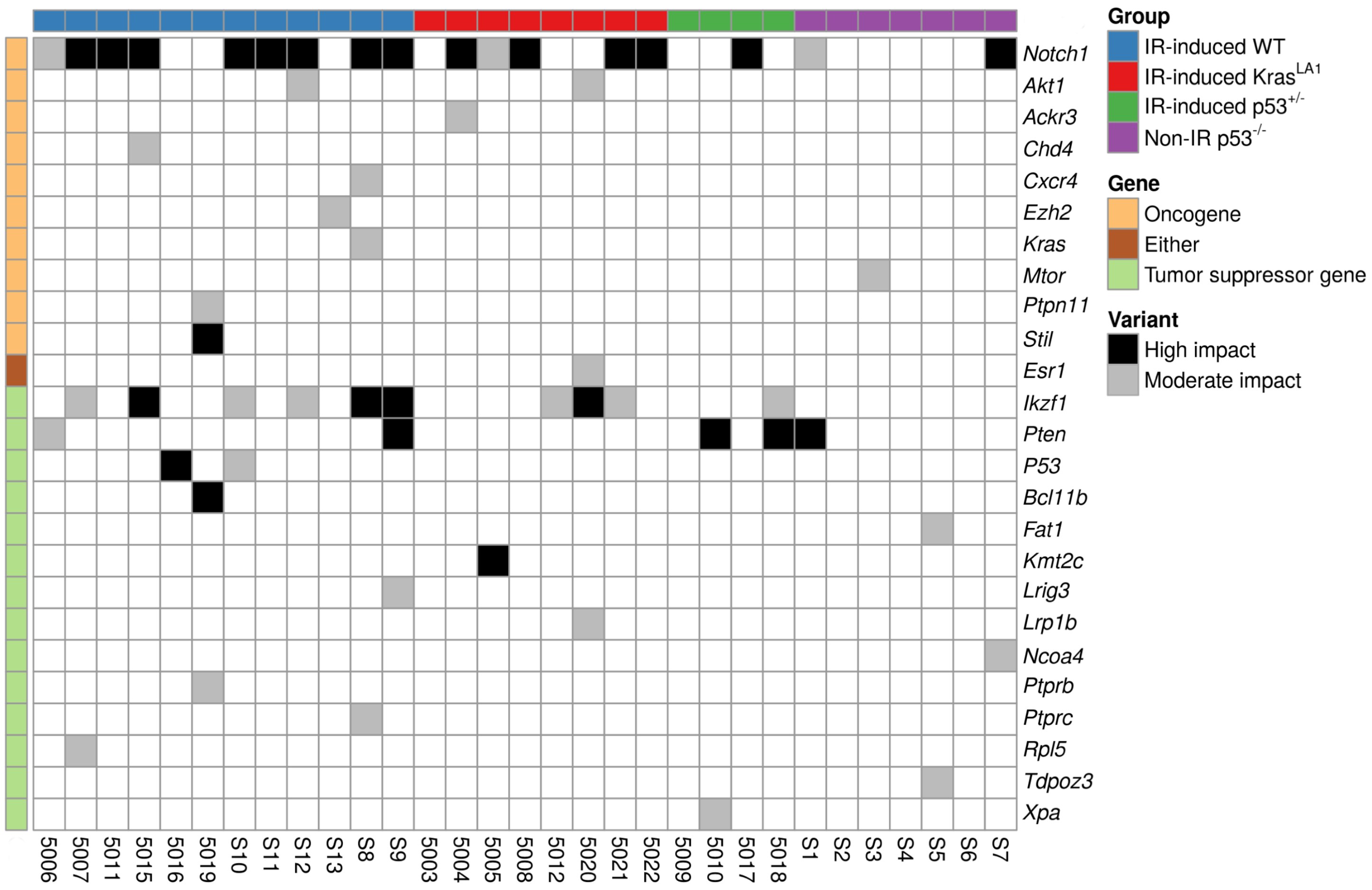


**Figure S4.** All nonsynonymous mutations in COSMIC genes across thymic lymphomas. Of note, 90% (18/20) of IR-induced WT and IR-induced Kras$^{LA1}$ lymphomas did not harbor mutations in p53. Among lymphomas that retained functional p53, approximately 83% (15/18) of these tumors harbored mutations *Notch1* and/or *Ikzf1*. Three p53 WT lymphomas that did not harbor mutations in either *Notch1* or *Ikzf1* are 5019 (mutations in *Ptpn11*, *Stil*, *Bcl11b* and *Ptprb*), S13 (mutation in *Ezh2*) and 5003 (mutation in *Kras*).

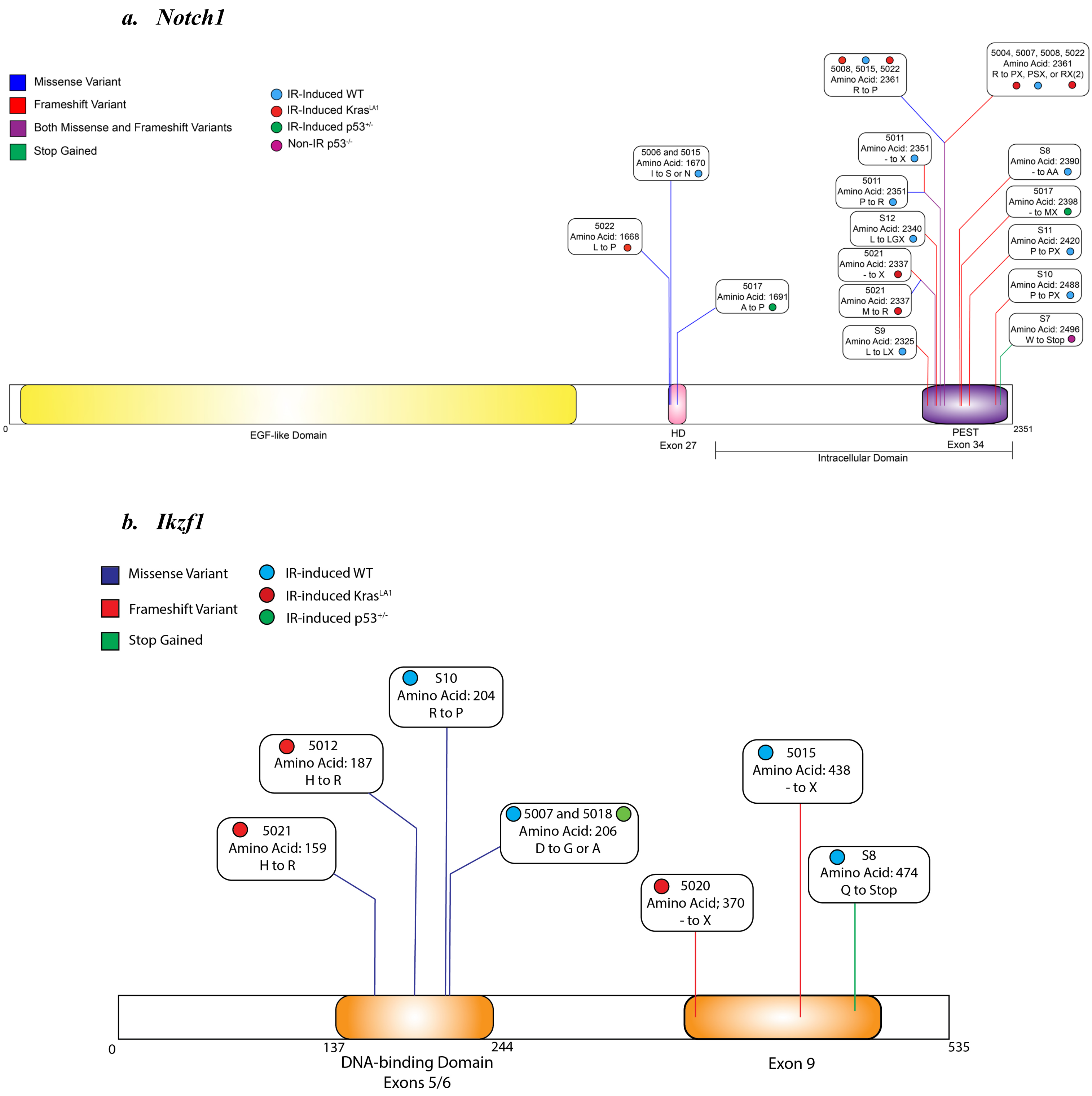


**Figure S5.** Schematic of nonsynonymous mutations in *Notch1* and *Ikzf*. All mutations listed in the plots were validated by Sanger sequencing. Colors of each dot represent lymphomas from mice with different genotypes. Colors of each line represent different types of somatic mutations. HD: heterodimerization domain; PEST: proline, glutamic acid, serine, threonine-rich domain

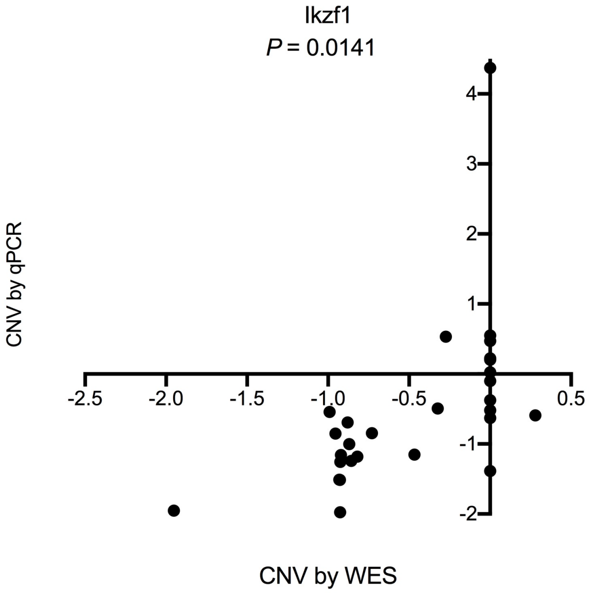


**Figure S6**. Validation of copy number variations (CNV) of *Ikzf1* determined by whole exome sequencing (WES) using qPCR. qPCR was performed using the same genomic DNA used for WES. *P* value was calculated by Kendall's W test for concordance, corrected for ties.

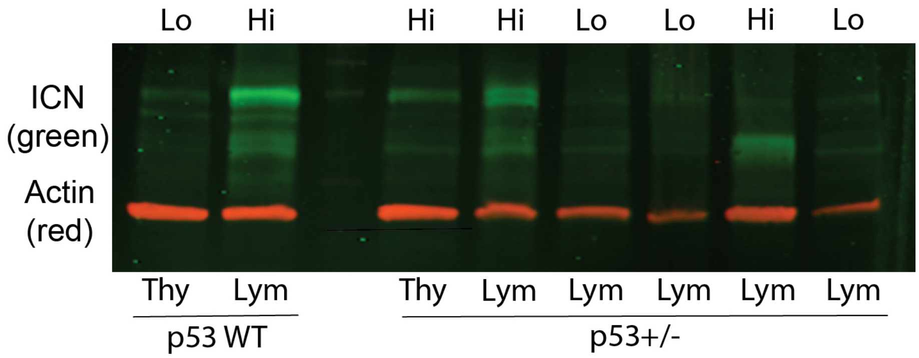


**Figure S7.** Representative Western blot that detects the expression of intracellular Notch1 (ICN) protein. The expression of ICN (green) was defined as either low (Lo) or high (Hi). Of note, we detected ICN protein with different molecule weight, which is likely due to different truncated mutations that occurred in the PEST domain of Notch1. Actin (red) was used as a control for protein loading. Thy: normal thymus; Lym: thymic lymphoma.

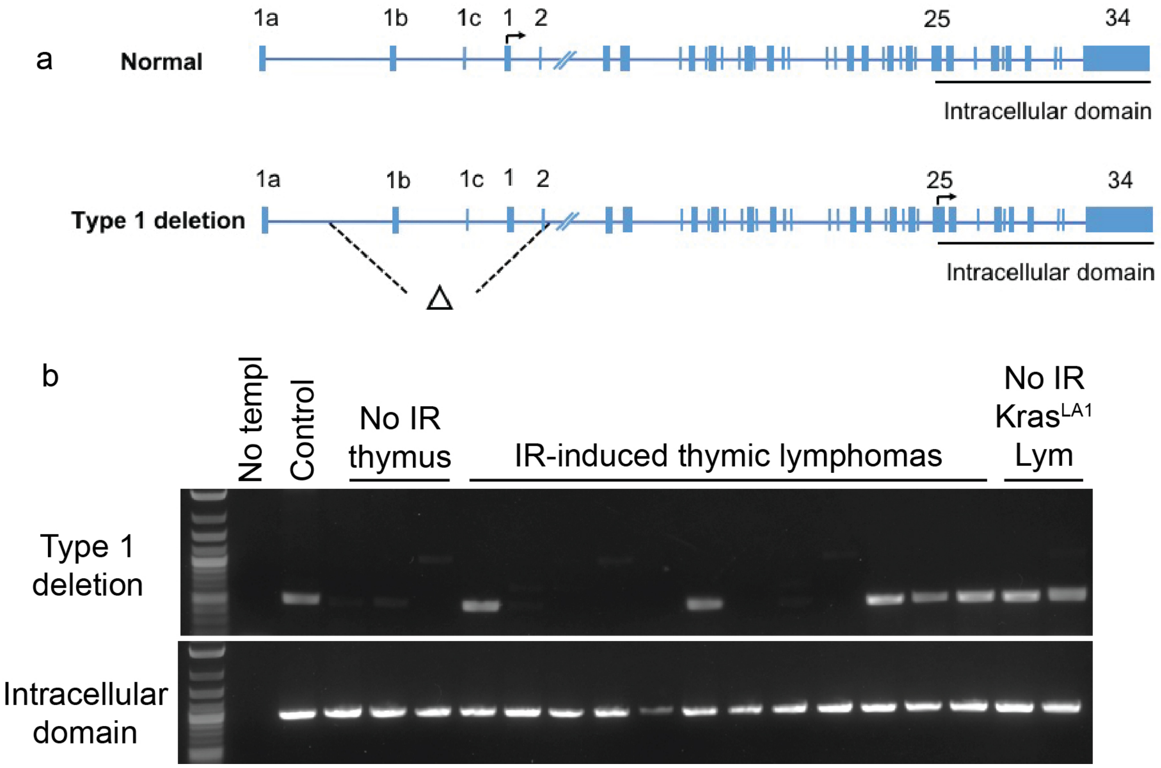


**Figure S8.** Detection of type I deletion of *Notch1*. **a,** Schematics of type 1 deletion of *Notch1*. Type 1 deletion removes the 5′ proximal promoter and exon 1 of *Notch1*, which activates transcription from the cryptic promoter in exon 25. **b,** Representative image showing the detection of *Notch1* type 1 deletion by PCR. Type 1 deletion was detected in radiation-induced thymic lymphomas across various genotypes as well as in thymic lymphomas from unirradiated Kras$^{LA1}$ mice. Amplification of a region inside the intracellular domain was used as a positive control for the presence of genomic DNA.

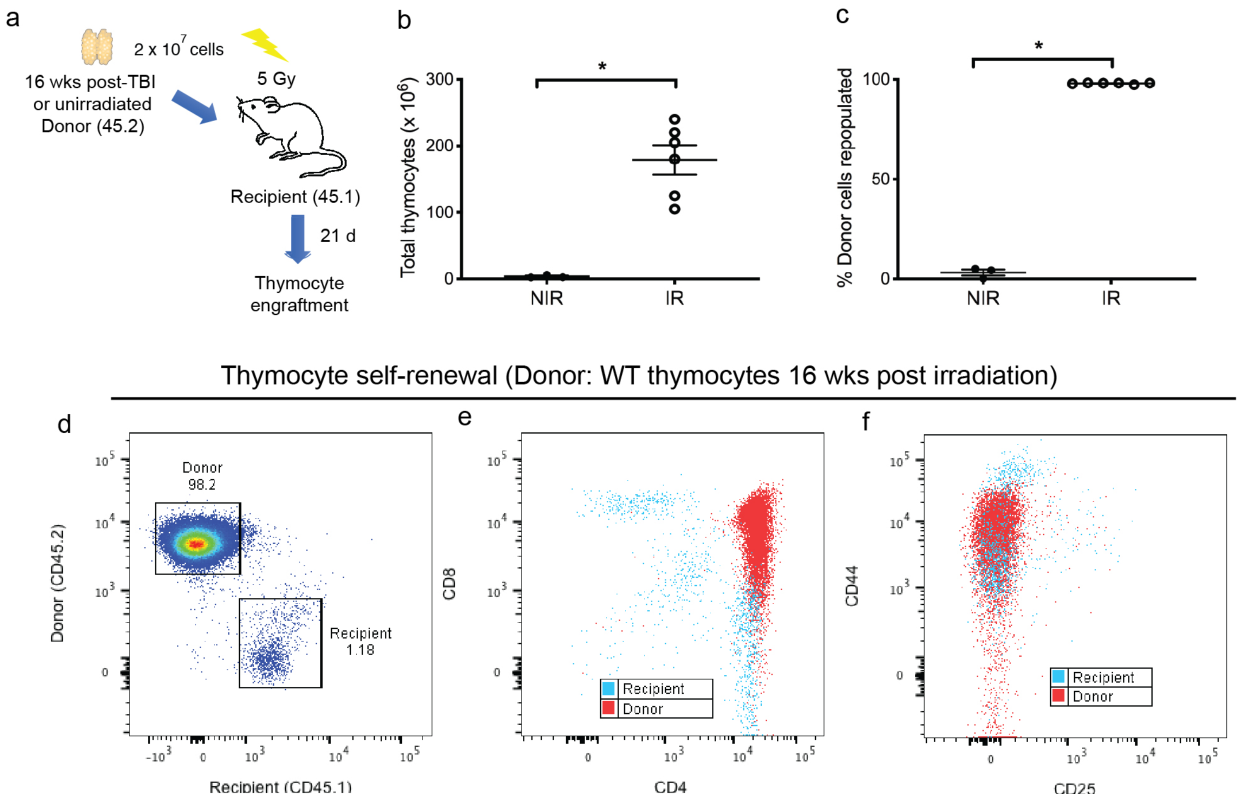


**Figure S9.** Self-renewal of thymocytes harvested from wild-type mice 16 weeks post-irradiation. **a,** 2 x $10^7$ donor thymocytes were harvested from age matched C57BL/6J mice (CD45.2) either 16 weeks after 1.8 Gy x 4 TBI (IR) or unirradiated (NIR). Donor thymocytes were injected intravenously into B6.SJL recipients (CD45.1) 6 hours after 5 Gy TBI. Thymocytes were harvested from recipients 21 days after transplantation. **b and c,** The number of total thymocytes (**b**) and the engraftment of donor thymocytes (**c**) were assessed in recipients that received thymocytes from donors that were either unirradiated (NIR, n=3) or previously treated with TBI (IR, n=6). Data are presented as mean ± SE. *$P$<0.05 by Mann-Whitney U test. **d-f,** Self-renewal of thymocytes harvested from wild-type mice 16 weeks post-irradiation. Representative flow cytometry plots that distinguish thymocytes derived from donor (CD45.2) vs. recipients (CD45.1). Of note, the majority of donor-derived thymocytes capable of self-renewal were $CD4^{+}CD8^{+}$ and also positive for CD44.

| Primer Name | Forward Sequence | Reverse Sequence |
|---|---|---|
| Ikzf1E5.3 | CAGGAGTTGGAGGCATTCGA | GGAAAACTCCCAGGCCTTACC |
| Ikzf1E6.2 | CAGTGTGGGGCCTCCTTTAC | GAGTGCGTCCTCAGGTGG |
| Ikzf1E8 | AGGAAGAAACTAACCACAACGAGA | AGCTCTTACGTTTGGCGACA |
| Ikzf1E9.2 | GCCAGTCATCAGCTCCATGT | TCAAAGGGATCCCGAAAGCC |
| Notch1E26 | TATACCTCCTCAGCCCCCTG | GGGTACATGCTCAGCACAGT |
| Notch1E27.2 | ACTGTGCCATGTGTGTTCTC | GACGCAAGAGCACCTAGGAA |
| Notch1E34.1 | GCTCCCTCATGTACCTCCTG | ATAGCATGATGGGGCCACTA |
| Notch1E34.2 | ATAGCATGATGGGGCCACTA | TTTTCCTGGTCAGGGTGAAG |
| Notch1E34.3 | ACTTGTCAGATGTGGCCTCG | GCAGCCACAGAACTTACAGC |
| Notch1E34.4 | CTCCCCATTCCAGCAGTCTC | AGGTGCAGCCACAGAACTTA |
| Notch1E34.5 | GCACGGGGGCTATGAATTTC | CACATCACTGCCATCCTCCA |
| PtenE5 | AGACCATAACCCACCACAGC | CAGCTTACCTTTTTGTCTCTGG |
| PtenE7 | GGTCTGCCAGCTAAAGGTGA | CAATAGAACTGAATTGCAAACCT |
| PtenE8 | ACCAGGACCAGAGGAAACCT | TATCGGTTGGCCTTGTCTTT |
| TRP53E7 | ACCTGGATCCTGTGTCTTCC | CTACCACGCGCCTTCCTAC |

**Table S1.** Summary of primers used for Sanger sequencing

| Pathway | Oncogene/Tumor Supressor | Type of Alteration | WT (n=12) | KrasLA1 (n=8) | p53+/- (n=4) | NIR p53 -/- (n=7) |
|---|---|---|---|---|---|---|
| Notch Signaling | | | | | | |
| Notch1 | Oncogene | Mutation | 75.0% | 62.5% | 25.0% | 28.6% |
| Ikzf1 | Tumor Supressor | CNVs | 75.0% | 62.5% | 50.0% | 28.6% |
| Ikzf1 | Tumor Supressor | Mutation | 50.0% | 37.5% | 25.0% | 0.0% |
| PI3K/AKT/mTOR pathway | | | | | | |
| Akt1 | Oncogene | Mutation | 8.3% | 12.5% | 0.0% | 0.0% |
| Pten | Tumor Supressor | CNVs | 25.0% | 0.0% | 25.0% | 42.9% |
| Pten | Tumor Supressor | Mutation | 16.7% | 0.0% | 50.0% | 14.3% |
| Mtor | Oncogene | CNVs | 8.3% | 0.0% | 75.0% | 28.6% |
| Mtor | Oncogene | Mutation | 0.0% | 0.0% | 0.0% | 14.3% |
| MEK/ERK Pathway | | | | | | |
| Kras | Oncogene | Mutation | 8.3% | 100.0% | 0.0% | 0.0% |
| Flt3 | Oncogene | CNVs | 0.0% | 0.0% | 25.0% | 0.0% |
| Ptpn11 | Oncogene | Mutation | 8.3% | 0.0% | 0.0% | 0.0% |
| Cxcr4 | Oncogene | Mutation | 8.3% | 0.0% | 0.0% | 0.0% |
| Ackr3 | Oncogene | Mutation | 0.0% | 12.5% | 0.0% | 0.0% |
| Epigenetic Modifiers | | | | | | |
| Chd4 | Oncogene | Mutation | 8.3% | 0.0% | 0.0% | 0.0% |
| Dnmt3a | Tumor Supressor | CNVs | 0.0% | 0.0% | 25.0% | 42.9% |
| Esr1 | Oncogene | Mutation | 0.0% | 12.5% | 0.0% | 0.0% |
| Bcl11b | Tumor Supressor | CNVs | 50.0% | 50.0% | 75.0% | 28.6% |
| Bcl11b | Tumor Supressor | Mutation | 8.3% | 0.0% | 0.0% | 0.0% |
| Dicer1 | Tumor Supressor | CNVs | 66.7% | 75.0% | 50.0% | 71.4% |
| Ezh2 | Oncogene | Mutation | 8.3% | 0.0% | 0.0% | 0.0% |
| Kmt2c | Tumor Supressor | Mutation | 0.0% | 12.5% | 0.0% | 0.0% |
| Hippo Pathway | | | | | | |
| Nf2 | Tumor Supressor | CNVs | 50.0% | 62.5% | 50.0% | 28.6% |
| Fat1 | Tumor Supressor | Mutation | 0.0% | 0.0% | 0.0% | 14.3% |
| Wnt Signaling | | | | | | |
| Lrig3 | Tumor Supressor | CNVs | 25.0% | 0.0% | 25.0% | 57.1% |
| Lrig3 | Tumor Supressor | Mutation | 8.3% | 0.0% | 0.0% | 0.0% |
| Cdh11 | Tumor Supressor | CNVs | 50.0% | 0.0% | 0.0% | 100.0% |
| Rspo3 | Oncogene | CNVs | 25.0% | 12.5% | 50.0% | 14.3% |
| Lrp1b | Tumor Supressor | Mutation | 0.0% | 12.5% | 0.0% | 0.0% |
| DNA Damage Response | | | | | | |
| p53 | Tumor Supressor | CNVs | 8.3% | 12.5% | 100.0% | 100.0% |
| p53 | Tumor Supressor | Mutation | 16.7% | 0.0% | 0.0% | 0.0% |
| Fancg | Tumor Supressor | CNVs | 58.3% | 37.5% | 100.0% | 57.1% |
| Rpl5 | Tumor Supressor | Mutation | 8.3% | 0.0% | 0.0% | 0.0% |
| Xpa | Tumor Supressor | Mutation | 0.0% | 0.0% | 25.0% | 0.0% |
| Others | | | | | | |
| Ncoa4 | Tumor Supressor | Mutation | 0.0% | 0.0% | 0.0% | 14.3% |
| Stil | Oncogene | Mutation | 8.3% | 0.0% | 0.0% | 0.0% |
| Pold1 | Tumor Supressor | CNVs | 33.3% | 62.5% | 75.0% | 14.3% |
| Ptprb | Tumor Supressor | Mutation | 8.3% | 0.0% | 0.0% | 0.0% |
| Ptprc | Tumor Supressor | Mutation | 8.3% | 0.0% | 0.0% | 0.0% |
| Tdpoz5 | Tumor Supressor | CNVs | 16.7% | 0.0% | 0.0% | 71.4% |
| Tdpoz3 | Tumor Supressor | Mutation | 0.0% | 0.0% | 0.0% | 14.3% |

**Table S2.** Summary of mutations and copy number variations in COSMIC genes among different cohorts of thymic lymphomas. Note: the frequency of Kras mutation and p53 mutation in $Kras^{LA1}$ and p53 deficient mice, respectively, is defined as 100%.

**Table S3.** Summary of validated codon change in COSMIC genes

| Gene | Sample | Treatment Group | Exon | Location | Consequence | cDNA position | Protein position | Codon Change | Sanger Codon Change |
|---|---|---|---|---|---|---|---|---|---|
| Akt1 | 5020 | IR-induced KrasLA1 | 4 | 12:112659599 | missense_variant | 608 | 80 | Tgg/Cgg | gtgg/gCtgg |
| Ikzf1 | 5021 | IR-induced KrasLA1 | 5 | 11:11748566 | missense_variant | 1034 | 159 | cAt/cGt | cat/cGt |
| Ikzf1 | 5012 | IR-induced KrasLA1 | 6 | 11:11754085 | missense_variant | 1118 | 187 | cAc/cGc | cac/cGc |
| Ikzf1 | 5020 | IR-induced KrasLA1 | 9 | 11:11769080-11769082 | frameshift_variant | 1665-1666 | 369-370 | -/C | cccccc/cccccccC |
| Notch1 | 5022 | IR-induced KrasLA1 | 27 | 2:26466601 | missense_variant | 5267 | 1668 | cTg/cCg | ctg/cCg |
| Notch1 | 5021 | IR-induced KrasLA1 | 34 | 2:26460117 | missense_variant | 7274 | 2337 | aTg/aGg | atggggc/aGgAggG |
| Notch1 | 5004 | IR-induced KrasLA1 | 34 | 2:26459764 | stop_gained | 7627 | 2455 | Cag/Tag | cag/cCg |
| Notch1 | 5022 | IR-induced KrasLA1 | 34 | 2:26460045 | missense_variant | 7346 | 2361 | cGg/cCg | cgg/cCg |
| Notch1 | 5004 | IR-induced KrasLA1 | 34 | 2:26460045-26460046 | frameshift_variant | 7345-7346 | 2361 | cgg/cCTgg | cgg/cCTgg |
| Notch1 | 5008 | IR-induced KrasLA1 | 34 | 2:26460044-26460045 | frameshift_variant | 7346-7347 | 2361 | cgg/cgCCg | cggc/cCCc |
| Notch1 | 5008 | IR-induced KrasLA1 | 34 | 2:26460045 | missense_variant | 7346 | 2361 | cGg/cCg | cggc/cCCc |
| Notch1 | 5022 | IR-induced KrasLA1 | 34 | 2:26460044-26460045 | frameshift_variant | 7346-7347 | 2361 | cgg/cgCg | cggctg/cCGGCT |
| Notch1 | 5005 | IR-induced KrasLA1 | 34 | 2:26460538-26460549 | inframe_deletion | 6842-6853 | 2193-2197 | cTCGAGTCACCCCat/cat | ctcgagtcaccccat/cCTgagtcCTccc |
| Notch1 | 5021 | IR-induced KrasLA1 | 34 | 2:26460115-26460116 | frameshift_variant | 7275-7276 | 2337-2338 | -/A | tggg/tgAgg |
| Ikzf1 | 5018 | IR-induced p53 +/- | 6 | 11:11754142 | missense_variant | 1175 | 206 | gAc/gCc | gac/gCc |
| Pten | 5018 | IR-induced p53 +/- | 7 | 19:3281578 | frameshift_variant | 1744 | 266 | Aaa/aa | aaaaag/aaaag |
| Notch1 | 5017 | IR-induced p53 +/- | 27 | 2:26466533 | missense_variant | 5335 | 1691 | Gcc/Ccc | gcc/Ccc |
| Notch1 | 5017 | IR-induced p53 +/- | 34 | 2:26459932-26459933 | frameshift_variant | 7458-7459 | 2398-2399 | -/ATGGG | gca/ATG |
| Ikzf1 | 4556-S10 | IR-induced p53 WT | 6 | 11:11754136 | missense_variant | 1169 | 204 | cGg/cCg | cgg/cCC |
| Ikzf1 | 4556-S10 | IR-induced p53 WT | 6 | 11:11754137 | synonymous_variant | 1170 | 204 | cgG/cgC | cgg/cCC |
| Ikzf1 | 4556-S12 | IR-induced p53 WT | 6 | 11:11754142 | missense_variant | 1175 | 206 | gAc/gCc | gac/gCc |
| Ikzf1 | 5007 | IR-induced p53 WT | 6 | 11:11754144 | missense_variant | 1175 | 206 | gAc/gGc | gac/gGc |
| p53 | 4556-S10 | IR-induced p53 WT | 7 | 11:69589183 | missense_variant | 863 | 236 | Aat/Gat | aat/Gat |
| Ikzf1 | 4556-S9 | IR-induced p53 WT | 8 | 11:11761253-11761254 | frameshift_variant | 1365-1366 | 269-270 | -/G | gaa/gGaa |
| Pten | 4556-S9 | IR-induced p53 WT | 8 | 19:32817896-32817897 | frameshift_variant | 1810-1811 | 288 | gaa/gAaa | gaaaaag/gaaaaaAg |
| Ikzf1 | 4556-S8 | IR-induced p53 WT | 9 | 11:11769393 | stop_gained | 1978 | 474 | Cag/Tag | cag/Tag |
| Ikzf1 | 5015 | IR-induced p53 WT | 9 | 11:11769287-11769288 | frameshift_variant | 1872-1873 | 438-439 | -/CC | cgc/cCCgc |
| Notch1 | 5015 | IR-induced p53 WT | 27 | 2:26466595 | missense_variant | 5273 | 1670 | aTc/aAc | atc/aAc |
| Notch1 | 5006 | IR-induced p53 WT | 27 | 2:26466595 | missense_variant | 5273 | 1670 | aTc/aGc | atc/atGc |
| Notch1 | 4556-S8 | IR-induced p53 WT | 34 | 2:26459956-26459957 | frameshift_variant | 7434-7435 | 2390-2391 | -/AA | cag/caAAg |
| Notch1 | 4556-S10 | IR-induced p53 WT | 34 | 2:26459663-26459664 | frameshift_variant | 7727-7728 | 2488 | cca/ccCa | cca/ccCa |
| Notch1 | 5011 | IR-induced p53 WT | 34 | 2:26460075 | missense_variant | 7316 | 2351 | cCg/cGg | ccg/cGg |
| Notch1 | 4556-S11 | IR-induced p53 WT | 34 | 2:26459867-26459868 | frameshift_variant | 7523-7524 | 2420 | ccg/ccCg | ccgc/ccgG |
| Notch1 | 5007 | IR-induced p53 WT | 34 | 2:26460045-26460046 | frameshift_variant | 7345-7346 | 2361 | cgg/cCCTCgg | cgg/cCCTTgg |
| Notch1 | 5015 | IR-induced p53 WT | 34 | 2:26460045 | missense_variant | 7346 | 2361 | cGg/cCg | cggct/cCgAc |
| Notch1 | 4556-S12 | IR-induced p53 WT | 34 | 2:26460107-26460108 | frameshift_variant | 7283-7284 | 2340 | cta/ctGGGTa | cta/ctGGGTa |
| Notch1 | 4556-S9 | IR-induced p53 WT | 34 | 2:26460152-26460153 | frameshift_variant | 7238-7239 | 2325 | ctg/ctTg | ctg/ctTg |
| Notch1 | 5011 | IR-induced p53 WT | 34 | 2:26460073-26460074 | frameshift_variant | 7317-7318 | 2351-2352 | -/G | gat/gaAt |
| Notch1 | 4556-S7 | Non-IR p53 -/- | 34 | 2:26459640 | stop_gained | 7751 | 2496 | tGg/tAg | tgg/tAg |
